# Unlocking plant health survey data: an approach to quantify the sensitivity and specificity of visual inspections

**DOI:** 10.1101/2025.03.13.642969

**Authors:** Matt Combes, Nathan Brown, Robin N. Thompson, Alexander Mastin, Peter Crow, Stephen Parnell

## Abstract

Invasive plant pests and pathogens cause significant environmental and economic damage. Visual inspection remains a central tenet of plant health surveys, but its sensitivity (probability of correctly identifying the presence of a pest) and specificity (probability of correctly identifying the absence of a pest) is usually ignored. These parameters facilitate calculation of surveillance metrics which are critical for effective contingency planning and outbreak management. To address this, twenty-three citizen scientist surveyors assessed up to 175 oak trees for three symptoms of acute oak decline. The same trees were also assessed by an expert who has monitored these trees annually for over a decade. The sensitivity and specificity of surveyors was calculated using the expert data as the ‘gold-standard’. The utility of a workflow utilising Bayesian modelling was then examined using simulated data to estimate these parameters in the absence of a rarely available ‘gold-standard’ dataset. There was large variation in sensitivity and specificity between surveyors and symptoms, although the sensitivity was positively related to the number of symptoms on a tree. By leveraging surveyor observations of two symptoms from a minimum of 80 trees on two sites, with knowledge of whether a site has higher (∼0.6) or lower (∼0.3) true disease prevalence we show that sensitivity and specificity can be estimated without gold-standard data. We highlight that sensitivity and specificity will depend on the symptoms of a pest or disease, the individual surveyor, and the survey protocol. This has consequences for how surveys are designed to detect and monitor outbreaks, as well as the interpretation of survey data that is used to inform outbreak management.

**Author summary:** The increasing occurrence of emerging plant pests and diseases is affecting both agricultural and natural ecosystems. Effective management and control of such pests and diseases is much easier when they are detected early. Currently, visual surveys underpin plant health surveillance, but basic metrics of the reliability of visual detection such as the sensitivity (probability of correctly identifying a positive) and specificity (probability of correctly identifying a negative) are not routinely quantified. In this study, we first quantify the sensitivity and specificity of 23 trained citizen scientist surveyors at detecting three symptoms of acute oak decline, by comparing their symptom classifications against a dataset from an expert who has conducted long-term monitoring of these trees. We demonstrate how individuals vary greatly in their ability to detect symptoms, and how different symptoms are associated with different detection error. Secondly, based on this dataset we outline a workflow developed for scenarios realistic in plant health which utilises Bayesian modelling to estimate these parameters in the absence of a rarely available ‘gold-standard’ expert dataset. In summary, our results highlight variation in the reliability of visual detection, and we provide a workflow to calculate this and facilitate optimisation of risk-based surveillance strategies in plant health.

## Introduction

Biological invasions are a major cause of global ecosystem destabilisation, affecting both terrestrial and aquatic systems [1]. Such invasions are occurring with increasing frequency, and are projected to continue to increase towards 2050 [2], driven by factors such as human mediated transport, and those associated with environmental change (e.g. climate change and socio-economic activity) [3].

Invasive plant insect pests and pathogens (from hereon in collectively referred to as pests) are a pertinent example of the negative impacts of biological invasions. For example, *Xylella fastidiosa* was first detected in Europe in 2013, and identified as the causal agent of Olive Quick Decline Syndrome in Italy [4]. Genetic analyses revealed the bacterium was most likely introduced via ornamental coffee plants from Costa Rica [5], and is now projected to cost the industry in Italy alone up to ∼ € 5 billion [6].

As well as impacting agricultural systems, invasive plant pests can have a major impact on natural and semi-natural plant communities. *Hymenoscyphus fraxineus* is a fungus which is native to East Asia, and does not cause disease in its native range [7]. The fungus was introduced to Europe at least four decades ago [8], and is now causing mortality of European native *Fraxinus excelsior* and *Fraxinus angustifolia* trees across the continent, with *F. excelsior* even being deemed at risk of extinction in northern Europe [9].

Once the epidemic spread of such plant pests surpasses a certain threshold, eradication in the landscape may no longer be realistic [10]. Effective surveillance strategies are essential for early detection, and this maximises the likelihood of eradication [11]. Even if eradication is not possible, early detection enables early intervention strategies, which can reduce the overall costs and resources required for disease management [10]. Surveys for the detection of invading plant pests are usually led by visual inspection, with confirmation via molecular diagnostics[12, 13]. There is also an important role of citizen science in this process by increasing the number of potential reporters [14], which is highlighted by the discovery of *Dryocosmus kuriphilus* (oriental chestnut gall wasp) in the UK by a citizen scientist [15]. Although the sensitivity (probability of correctly identifying the presence of a pest) and specificity (probability of correctly identifying the absence of a pest) of laboratory tests are calculated [13], the probability of detecting a pest via initial visual inspection is routinely ignored. This makes it impossible to answer key practical questions from survey data, such as: what prevalence has an invader reached when it is first detected [16] or what is the probability that a pest is absent if it is not detected by a survey [12].

The lack of attention to the sensitivity and specificity of visual inspection is partly due to the rarity of ‘gold-standard’ reference datasets to calculate these parameters in the field. Therefore, approaches to enable estimation of sensitivity and specificity in the absence of a gold-standard are sorely needed. In this study, we aimed to address this knowledge gap surrounding sensitivity and specificity for visual plant health surveys. To achieve this, we used acute oak decline (AOD) as a case study due to the availability of study sites containing trees with known status for three different signs and symptoms (from hereon in collectively referred to as symptoms). We conducted experiments using citizen scientists to assess and record symptoms of AOD across two oak woodlands in southern England. The availability of an expert who has monitored these study sites annually in detail for over a decade provided the closest available to a ‘gold-standard’ expert dataset for visual inspection of symptoms. Next, we addressed the difficulty of calculating these parameters more widely across different plant pests and diseases by examining a workflow realistic for plant health to achieve this in the absence of a gold standard dataset. The specific objectives of this study were: (1) to provide a workflow to quantify sensitivity and specificity in the absence of a gold-standard dataset to facilitate optimisation of risk-based surveillance strategies in plant health; (2) to understand the extent to which sensitivity and specificity differ for visual inspection depending on the survey target (i.e. the symptom); (3) to understand to what extent surveyors differ in their sensitivity and specificity.

## Methods

### AOD as a case study

AOD is a decline disease involving patches of stem necrosis caused by a bacterial pathobiome and a native bupresid beetle that affects both UK native oak species, *Quercus robur* and *Q. petraea* [17–19]. The disease is characterised by the presence of four symptoms; externally visible weeping patches on oak stems, with underlying tissue necrosis in the sapwood, otherwise referred to as stem bleeds; the presence of cracks between bark plates with cavities of decayed tissue below; larval galleries of *Agrilus biguttatus* on more than 90% of affected trees, and externally visible D-shaped exit holes of the adult *A. bigutattus* on approximately a third of affected trees [17].

The presence of three externally visible characteristic symptoms (stem bleeds, bark cracks, *A. biguttatus* exit holes) (Fig 1), and the expert knowledge of AOD on the study sites enabled the design of surveys that could be used to calculate the sensitivity and specificity of citizen scientists at visually detecting morphologically different tree health symptoms.

**Fig 1.**
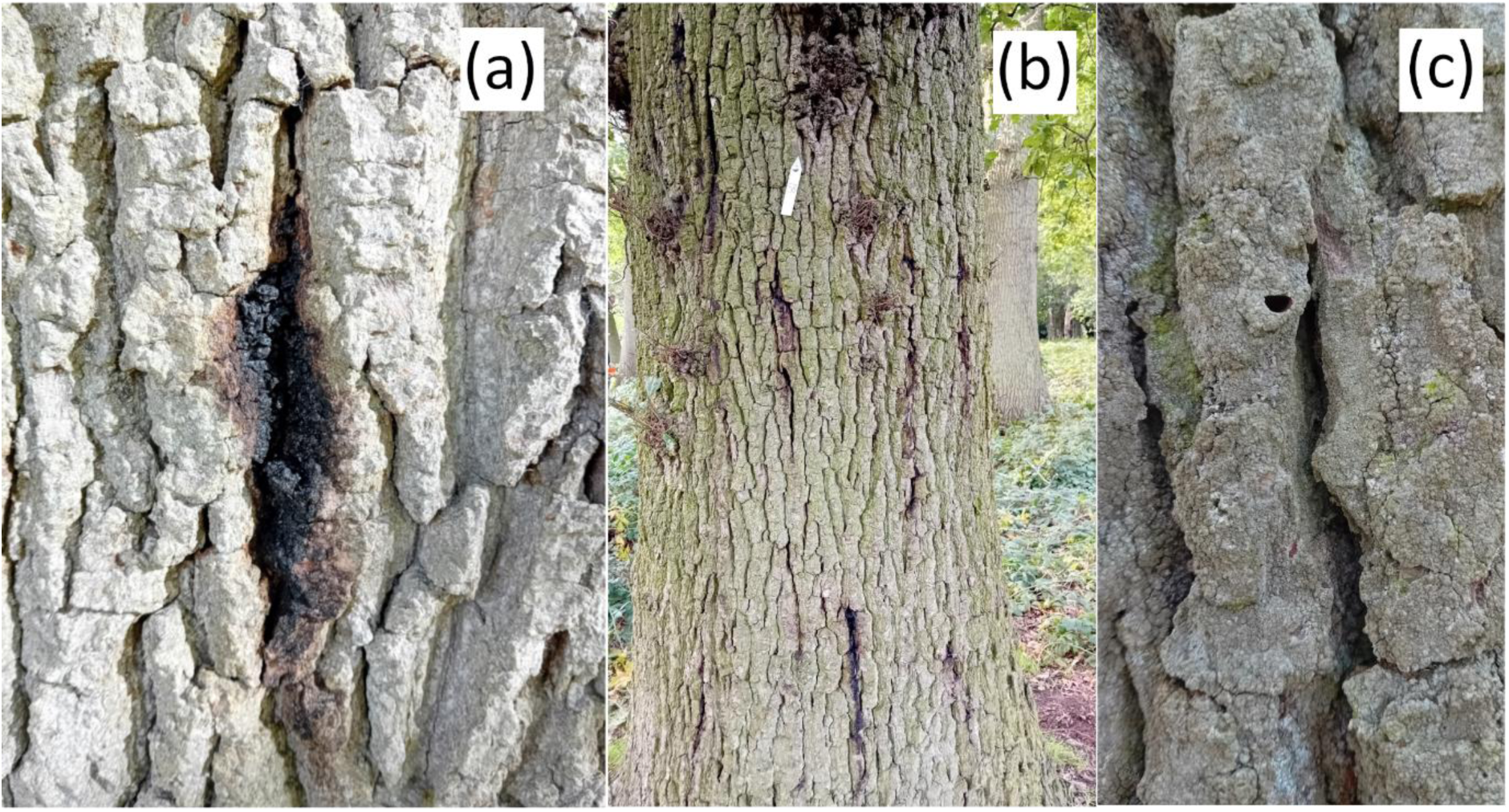
Three characteristic externally visible symptoms of AOD. [17]. (a) weeping patches on oak trunks/ stem bleeds; (b) AOD affected tree with stem bleeds and bark cracking between bark plates; (c) *Agrilus biguttatus* D-shaped exit holes (approximately 3-5 mm width).

### AOD surveys

Richmond Park (OS grid reference: TQ203744) and Hatchlands Park (OS grid reference: TQ06517) (Fig 2) were selected for AOD surveys based on their good public accessibility for citizen scientist surveyors, and the presence of knowledge of tree health status from annual AOD monitoring by an expert for over a decade (2009 at Hatchlands Park; 2010 at Richmond Park [20]). Walking routes of ∼4 km were mapped on both sites, and oak trees along these routes were labelled and the expert assessed these trees for the presence of the three externally visible AOD symptoms (Fig 2) one week prior to the first citizen scientist survey. In total, this involved 75 oak trees at Hatchlands Park and 100 at Richmond Park.

**Fig 2.**
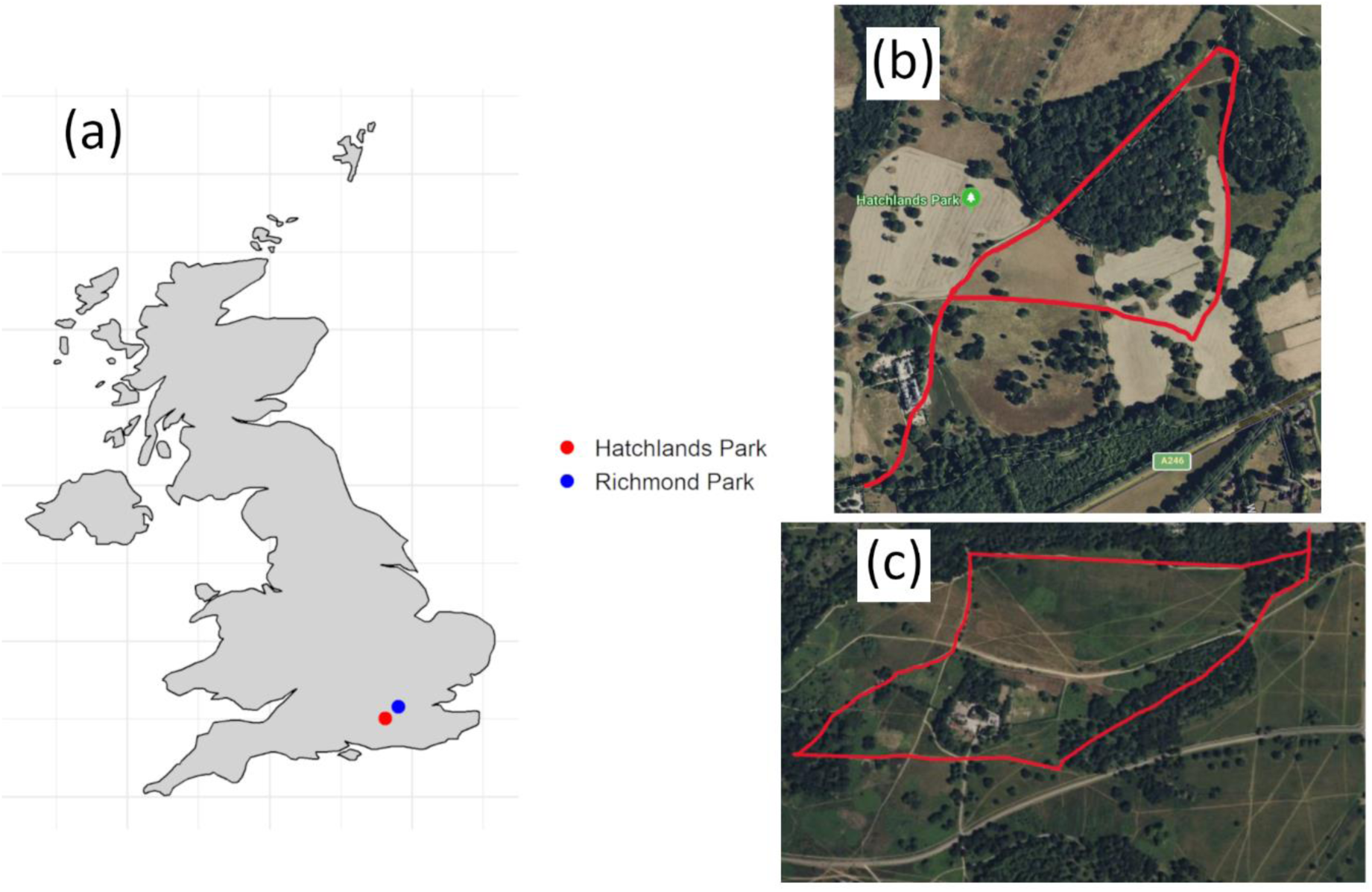
AOD survey sites. (a) in context of the UK, and ∼ 4 km routes at (b) Hatchlands Park and (c) Richmond Park where a combined total of up to 175 oak trees were assessed for three AOD symptoms by an expert and citizen scientist surveyors.

A total of 23 citizen scientists were recruited to conduct the survey, with only three of these surveyors not previously having undertaken either professional or volunteer tree work, and declaring no AOD knowledge. All citizen scientist surveyors received detailed AOD training prior to and on the survey days (S1 File). On the survey days (Richmond Park: 20/07/2022, 01/08/2022; Hatchlands Park: 21/07/2022, 02/08/2022), they were instructed to individually walk the specified ∼ 4 km routes and record the three external symptoms of AOD for each labelled tree (Fig 1). The expert assessor was present at each survey to note any changes in symptoms since their initial assessment.

### Ethics statement

Participants provided verbal consent to be part of the study, with all data analysed anonymously.

### Analyses of AOD survey data

The data on trees collected by citizen scientist surveyors was converted into presence and absence data for the three symptoms, and sensitivity and specificity of symptom detection was calculated using the expert assessor as a reference, or ‘gold-standard’.

The relationship between sensitivity, the type of symptom, the individual surveyor and the frequency of the symptoms were first visualised and then statistically analysed using generalised linear models with a binomial distribution and logit function. This was repeated to examine the relationship between specificity, the type of symptom and the individual surveyor. The significance of model parameters was assessed using type II likelihood ratio chi-square tests, with non-significant terms removed from models. Model interactions involving the individual surveyor and the frequency of the symptoms were not assessed due to a comparably limited dataset. The data were analysed using R software version 4.1.2 [21], with packages car [22], DHARMa [23], DescTools [24], and visualised using ggplot2 [25].

### Methods to quantify sensitivity and specificity in the absence of a ‘gold-standard’ expert dataset

Using maximum likelihood [26] and Bayesian approaches [27], the sensitivity and specificity of diagnostic tests can be estimated when two (or more) tests are applied to the same individuals across two (or more) populations with different true disease prevalences. These methods assume independence of diagnostic test sensitivity and specificity, although Dendukuri and Joseph [28] provide a methodology to account for non-independence. The application of these models for veterinary disease diagnostics using a Bayesian approach outlined below, is described in detail by Branscum et al. [29]. We apply these models to examine their utility for a workflow realistic for visual plant health inspections.

As described in Branscum et al. [29], the data for each tested individual can be separated into four categories: *y_11_* = test one is positive, test two is positive; *y_12_* = test one is positive, test two is negative; *y_21_* = test one is negative, test two is positive; *y_22_* = test one is negative, test two is negative. This count data from the four categories relates to the multinomial cell probabilities: *p_11_* = probability test one is positive, test two is positive; *p_12_* = probability test one is positive, test two is negative; *p_21_* = probability test one is negative, test two is positive; *p_22_* = probability test one is negative, test two is negative. For each population this can be characterised by Equation 1.

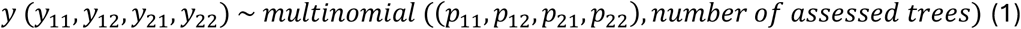

The multinomial cell probabilities can be expressed using eight parameters if covariance parameters (to account for non-independence of sensitivity and specificity between tests), are included in the model (Equation 2), and six if these covariance parameters are not included in the model (i.e. test sensitivity and specificity are independent) (Equation 3): True disease prevalence in population one, *π_1_*; True disease prevalence in population two, *π_2_*; Sensitivity of diagnostic test one, *Se_1_*; Sensitivity of diagnostic test two, *Se_2_*; Specificity of diagnostic test one, *Sp_1_*; Specificity of diagnostic test two, *Sp_2_*; Covariance between tests for disease positives, *CovD*^+^; Covariance between tests for disease negatives, *CovD*^−^.

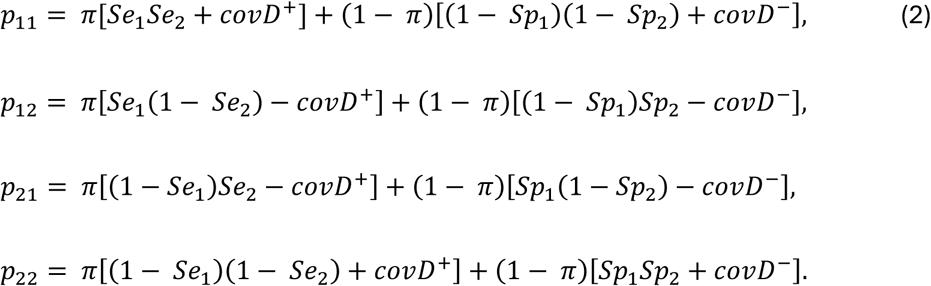

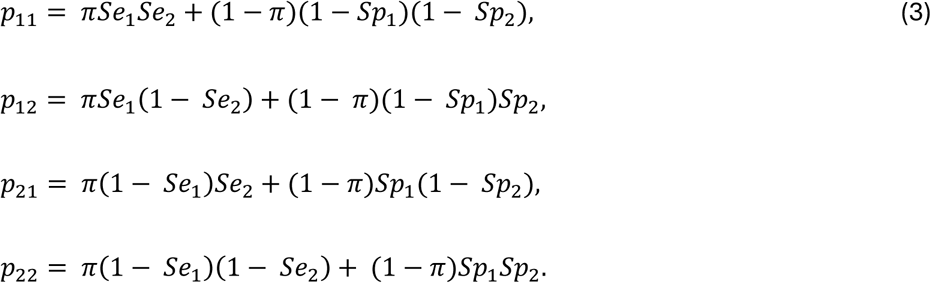

Markov chain Monte Carlo (MCMC) methods were used to iterate across a range of values for the parameters to estimate the probability distribution of observing the recorded data for the given value of the parameter, whilst accounting for the prior probability distribution of parameters [30]. This produces an estimate of the probability of a parameter value given the data (the posterior probability distribution) (Equation 4).

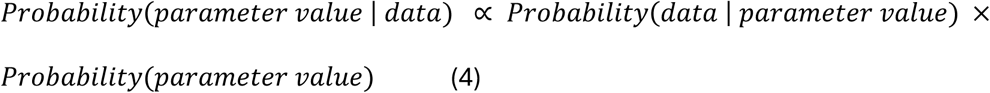

For this model, beta distributions were used to estimate the prior probability distributions for the true disease prevalence, and the test specificity and sensitivity [27, 29]. The priors for covariance parameters were modelled using a generalised beta distribution with uniform prior probability distributions, based on the minimum and maximum possible values [28, 29].

### A workflow to estimate sensitivity and specificity of visual plant health inspection in the absence of gold-standard data

Simulated survey datasets were used to explore field-realistic scenarios for a workflow for the quantification of the sensitivity and specificity of plant health visual inspection in the absence of a ‘gold-standard’ expert dataset. Sensitivity and specificity values for the visual detection of two symptoms were simulated based on the distributions of sensitivity and specificity for the visual detection of AOD stem bleed symptoms (Fig 3). To add non-independence between the detection of the two symptoms dependent on the disease status of a host (i.e. covariance between sensitivity and specificity of tests), we assumed a probability of 0.85 that symptom two would be scored as positive if symptom one was scored as positive and the host was disease positive, and that symptom two would be scored as negative if symptom one was scored as negative and the host was disease negative. This value was chosen in the absence of prior information on the relationship between sensitivity and specificity of symptom detection that can be expected in wider plant health, but was selected to test the robustness of a workflow using the covariance model, whilst also avoiding a more unrealistic scenario where the diagnostic result of two symptoms on a given plant are close to identical. The covariance values were calculated for simulated datasets (Equation 5), along with the minimum and maximum possible values for the covariance parameters given the simulated sensitivity and specificity values for surveyors (Equation 6) (S3 File).

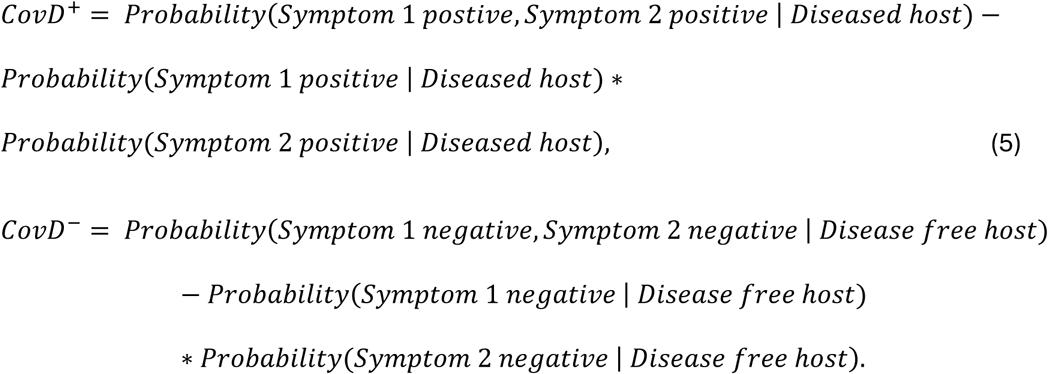

**Fig 3.**
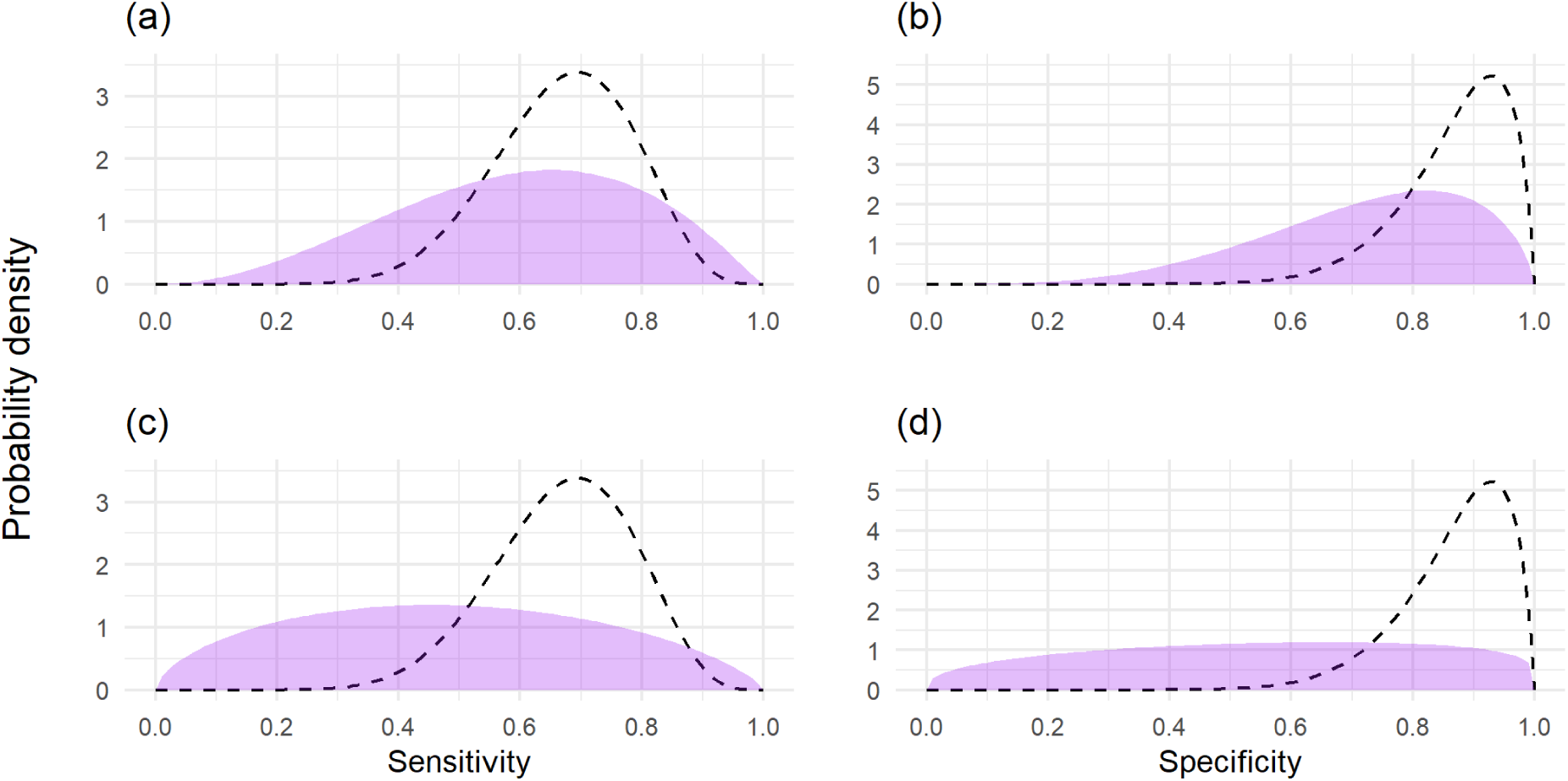
Prior probability distributions of sensitivity and specificity used in the simulated workflow, and the distributions used to simulate surveyor sensitivity and specificity. (a) Good prior knowledge of sensitivity, (b) good prior knowledge of specificity, (c) uninformed prior knowledge of sensitivity, (d) uninformed prior knowledge of specificity. The dashed line represents the distribution used to simulate surveyor sensitivity and specificity, and represents the distribution used for very good prior knowledge of sensitivity and specificity.

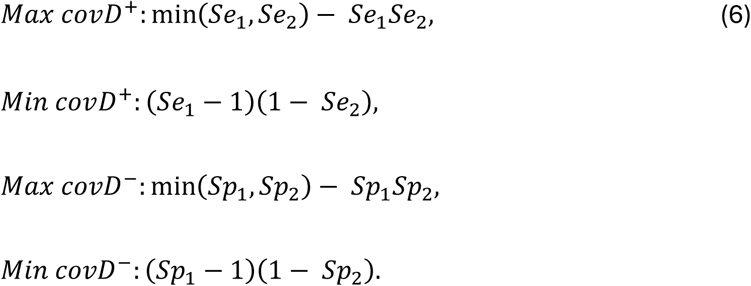

Based on experience with the AOD surveys, it was believed ∼25 surveyors can be organised to each inspect ∼100 trees in a day. We therefore tested a workflow using simulated datasets (either with or without covariance in the ability to detect each symptom) with the models above and 25 surveyors inspecting 80, 100 and 120 trees each from higher true disease prevalence (∼0.6), and lower true disease prevalence (∼0.3) areas (see S3 File for true disease prevalence values from each simulation). Surveyors then record the presence or absence of two disease symptoms on each assessed tree. This produces count data for *y_11_*, *y_12_*, *y_21_*, *y_22_*.

We anticipate informative prior knowledge of the higher true disease prevalence and the lower true disease prevalence areas (Fig 4; S3 File). Informative prior knowledge of these site locations can be reliably assumed from delimiting surveys that are conducted upon discovery of a pest [12]. We assume no prior knowledge of the covariance parameters between symptoms, but we anticipate knowledge of whether covariance may be present. This means we only include covariance parameters in the workflow when we simulate data with covariance. In the model, we specify uniform prior probability distributions between the minimum and maximum possible values (*cf.* [28, 29]) (Equation 6).

**Fig 4.**
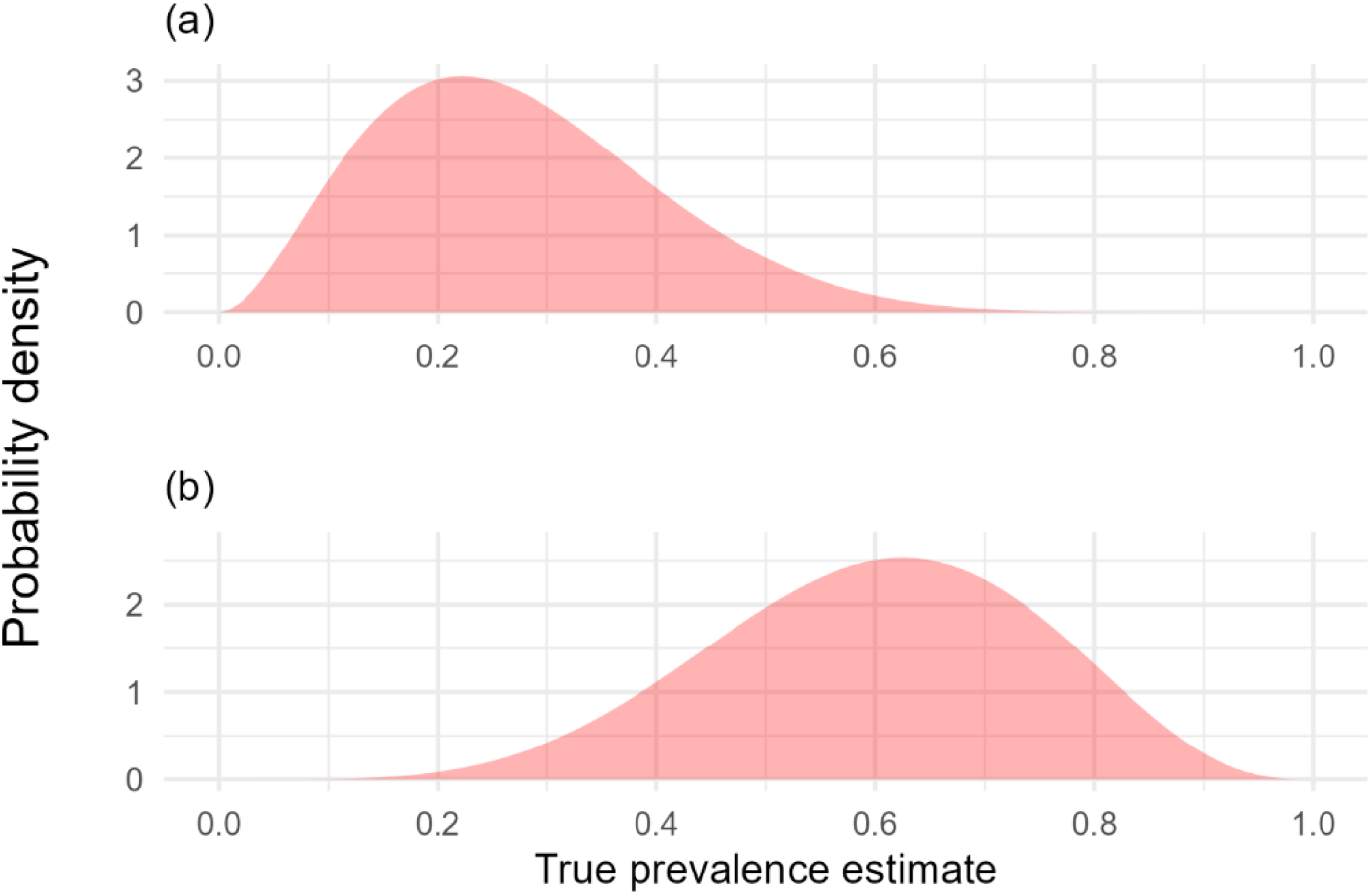
Prior probability distributions of (a) lower, (b) higher true disease prevalence locations used in the simulated workflow. Surveyor observations were simulated using 0.3 for lower true disease prevalence locations and 0.6 for higher true disease prevalence locations.

Three scenarios with different prior knowledge of the sensitivity and specificity for the two symptoms were tested, with the ability to estimate the sensitivity and specificity of symptom one assessed in subsequent analyses: (1) good prior knowledge of sensitivity and specificity of both symptoms; (2) poor prior knowledge of the sensitivity and specificity of symptom one, but very good prior knowledge of the sensitivity and specificity of symptom two; (3) poor prior knowledge of the sensitivity and specificity of both symptoms (Fig 3; S3 File). These scenarios were examined (1) excluding covariance parameters with a dataset without simulated covariance, and (2) including covariance parameters on a dataset with simulated covariance.

For each surveyor, MCMC methods (using Gibbs sampling) were then used to iterate across a range of values for each model parameter as described previously. A single chain had 10,000 burn-in iterations, 2000 adaptive iterations and 500,000 sample iterations. To check model convergence, this was repeated with nine additional chains and the potential scale reduction factor of the Gelman-Rubin statistic autocorrelation of the sample (psrf) was calculated, with values above 1.1 indicating inadequate model convergence. Results were excluded from further analyses if model convergence was inadequate.

Posterior estimates of the alpha and beta parameters for the beta distribution of a surveyor’s sensitivity and specificity were obtained for each iteration of the chain. These were then each thinned at every 10^th^ value after the initial 12,000 iterations, providing 48,800 alpha and beta value estimates for the beta distributions of sensitivity and specificity. The sensitivity and specificity of an individual surveyor was then estimated by randomly sampling values from each of these 48,800 possible beta distributions. The 48,800 estimates of sensitivity and specificity for each surveyor were combined to provide a group-level dataset of up to 1,220,000 values to estimate the group-level distribution of these parameters (i.e. distribution surveyors were sampled from). Kernel density estimates were then computed, and the results were visualised using a density plot.

Bayesian modelling and analyses were performed using Just another Gibbs Sampler (JAGS) program version 4.3.0 [31] in R software version 4.1.2 [21] with the packages runjags [32], coda [33], epiR [34] and fitdistrplus [35], and visualised using the ggplot2 [25].

## Results

### AOD case study

Calculation of surveyor sensitivity and specificity of visual detection of symptoms using expert data as a gold-standard revealed large variation between the three AOD symptoms and between individual surveyors (Fig 5; S2 File). Sensitivity differed between symptoms, and between surveyors, but increased with the frequency of symptoms (Fig 6). Significant interactions were present between both the type of symptom and frequency of symptom (LR Chisq = 49.803, df = 2, p < 0.001), and between the type of symptom and the surveyor (LR Chisq = 106.58, df = 44, p < 0.001). This highlights that the positive relationship between symptom frequency and sensitivity differs between symptoms, and that surveyor performance was not consistent relative to the symptom. Specificity was higher than sensitivity for 22 of 23 surveyors, but also differed between symptoms, and surveyors, with a significant interaction present between the two (LR Chisq = 192.19, df = 44, p < 0.001).

**Fig 5.**
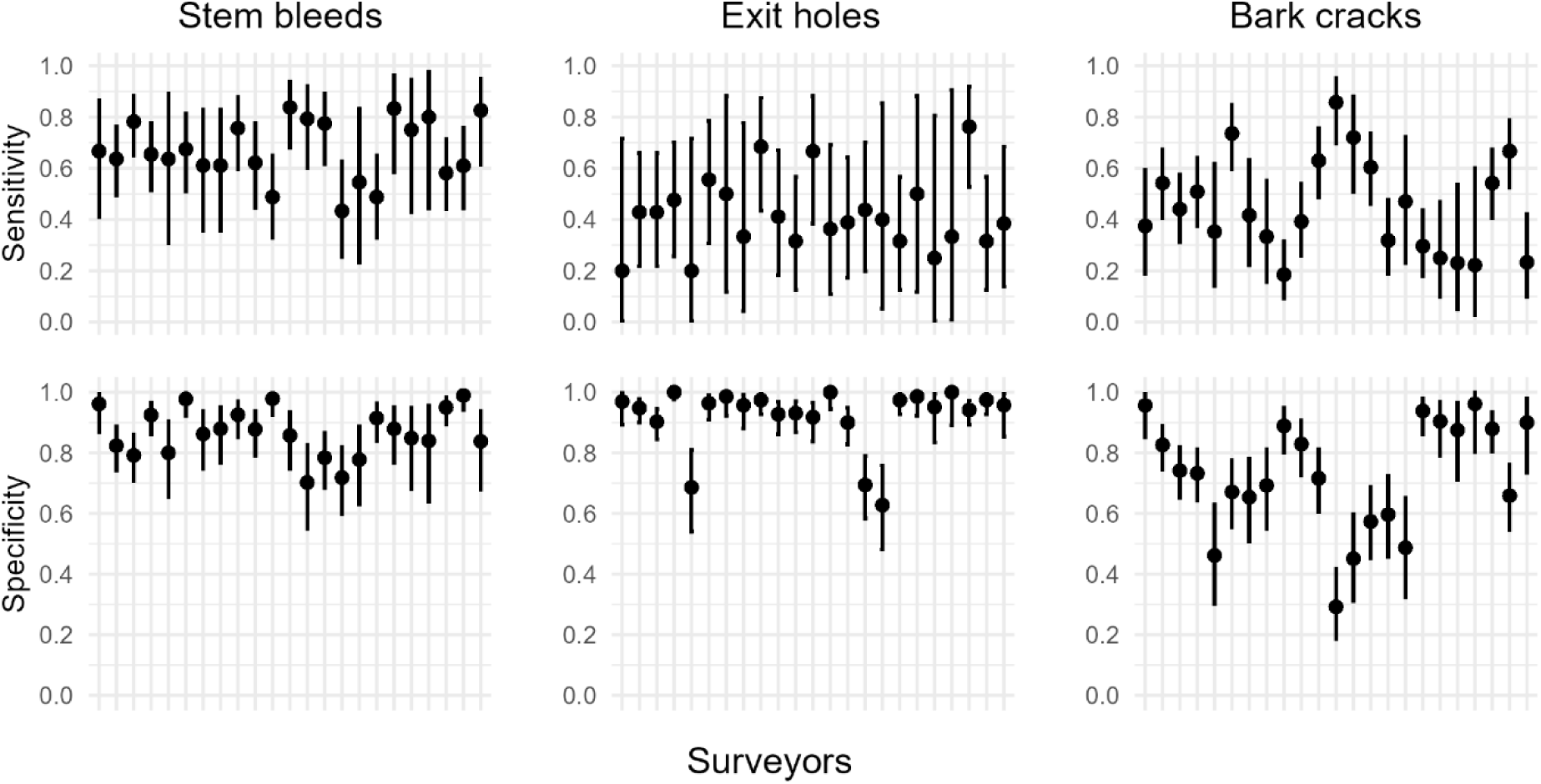
Sensitivity and specificity of three externally visible AOD symptoms on the main stem. The presence/ absence of bleeds/ weeping patches, *Agrilus* beetle exit holes, cracks between bark plates was assessed by 23 surveyors on up to 175 oak trees. Values were calculated using expert data as a reference standard, with error bars representing 95% confidence intervals. Surveyors are ordered consistently between plots on the x axes.

**Fig 6.**
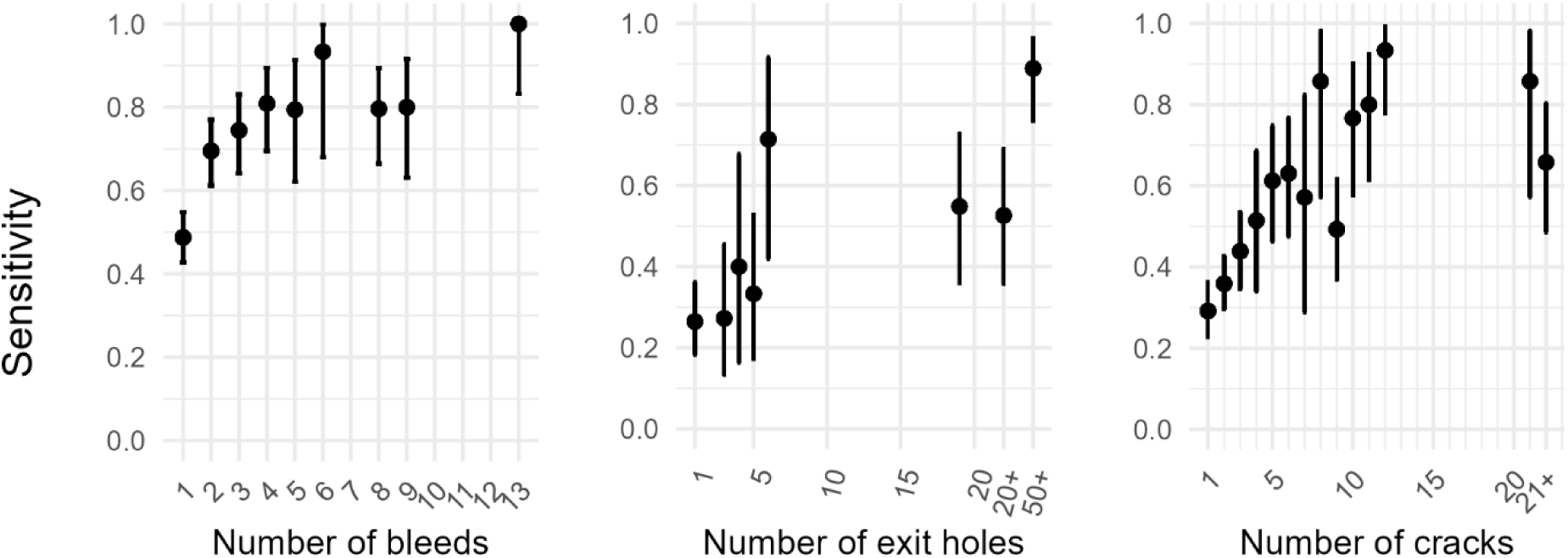
Sensitivity of three externally visible AOD symptoms on the main stem against the frequency of each symptom on a host. The presence/ absence of bleeds/ weeping patches, *Agrilus* beetle exit holes, cracks between bark plates was assessed by 23 surveyors on up to 175 oak trees. Data was pooled for surveyors, sensitivity was calculated using expert data as a reference standard, and then compared against the number of symptoms scored on a tree from the expert data. Error bars represent 95% confidence intervals. Values on the x axes ‘20+’, ‘50+’, ‘21+’ represent categorical data that were recorded at a lower precision due to the high frequency of symptoms on the host.

Overall, stem bleeds were the most reliable symptom to assess when accounting for both sensitivity and specificity, whilst *A. bigutattus* beetle exit holes had a high specificity but a low sensitivity (i.e. it was difficult to detect a D-shaped 3-5 mm exit hole on a tree and not common to falsely report one). Bark cracks were the least reliable symptom to visually detect (Fig 5), with many individuals performing similar to random chance (S1 Fig).

### A workflow to estimate sensitivity and specificity in wider plant health

The workflow enabled sensitivity and specificity of surveyors to be estimated with reasonable accuracy when as few as 80 trees were surveyed in locations with only informative prior knowledge of whether locations had a ‘higher’ or ‘lower’ true disease prevalence (Fig 7; S3 File). A notable outlier is present in Fig 7b, and represents a simulation where true disease prevalence only differed by 0.1 between low and high true disease prevalence locations, and the simulated surveyor specificity value was not supported by the prior probability distribution. Indeed, model estimates for individual surveyors were less reliable when surveyor sensitivity and specificity for either symptom was at a tail of a prior distribution (S18 Fig; S3 File), and occasionally resulted in models not converging (see below).

**Fig 7.**
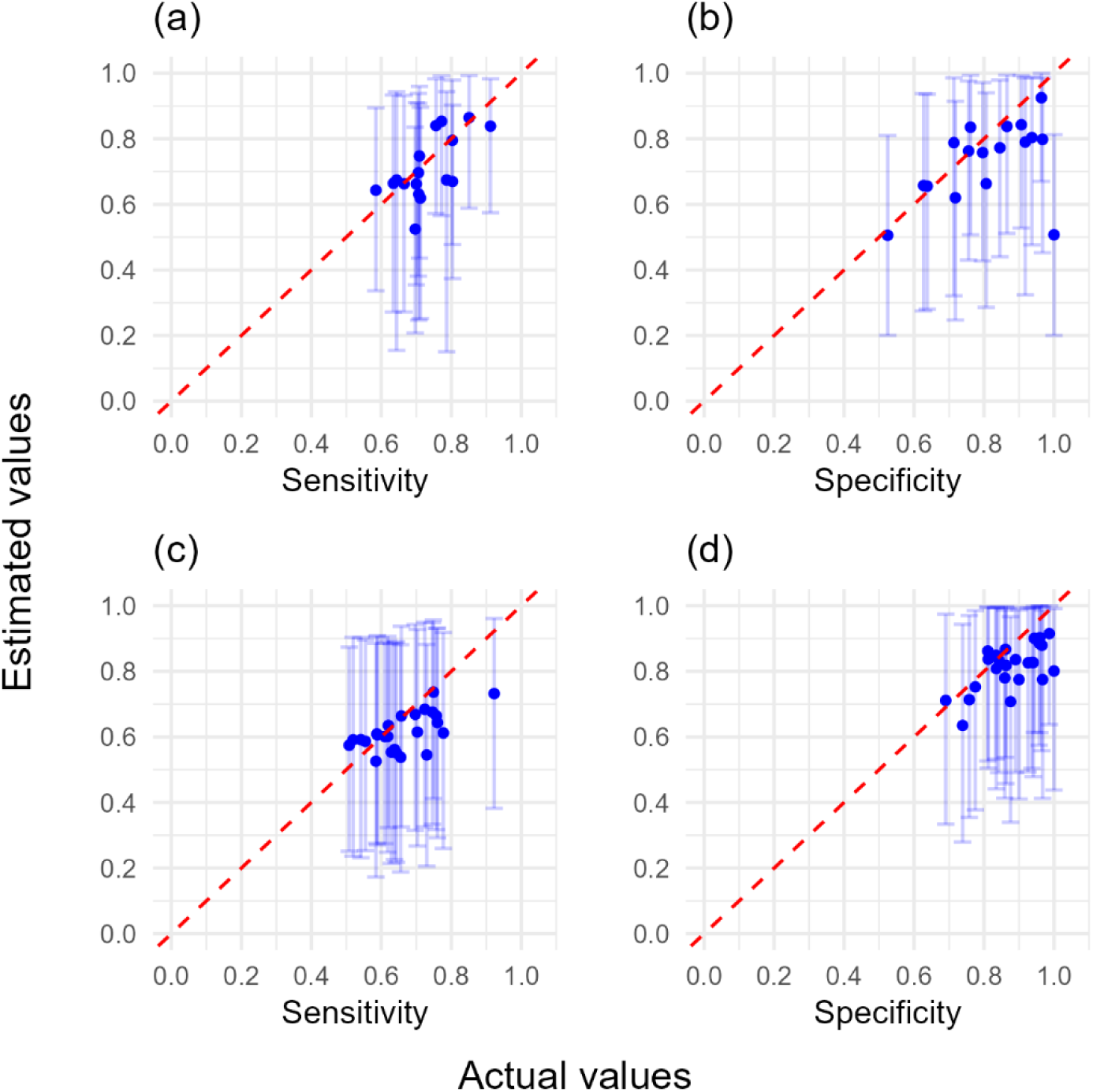
The workflow estimated sensitivity and specificity for simulated surveyors against actual sensitivity and specificity values. This workflow used poor sensitivity and specificity prior distributions with (a, b) no covariance model with no simulated covariance, (c, d) covariance model with simulated covariance. In both instances 80 trees each were assessed in higher and lower true disease prevalence locations. Error bars represent 95% confidence intervals, and dashed line represents perfect agreement between workflow estimates and actual values.

The group-level distributions of sensitivity and specificity the individual surveyors were simulated from could be estimated by combining sensitivity and specificity estimates from the 25 surveyors both in the presence and absence of covariance in the model and simulated data (Fig 8; S3 File). However, unsatisfactory model convergence (psrf > 1.1) occurred for some models without covariance parameters with poor prior information of sensitivity and specificity (80 trees 17/ 25 converged; 100 trees 20/25 converged; 120 trees 18/25 converged) (S3 File). This reduced surveyor numbers used in analyses and affected the accuracy of distribution estimates (Figs 8a and 8b). Informative prior information of the sensitivity and specificity of one, or both symptoms improved the accuracy of model outputs, although narrower ‘good’ and ‘very good’ prior knowledge of sensitivity and specificity did not necessarily improve individual estimates of these parameters because some surveyors were at the tails of these distributions. There was also minimal additional benefit in surveying either 100 or 120 trees per higher or lower true prevalence location (S2-17 Figs).

**Fig 8.**
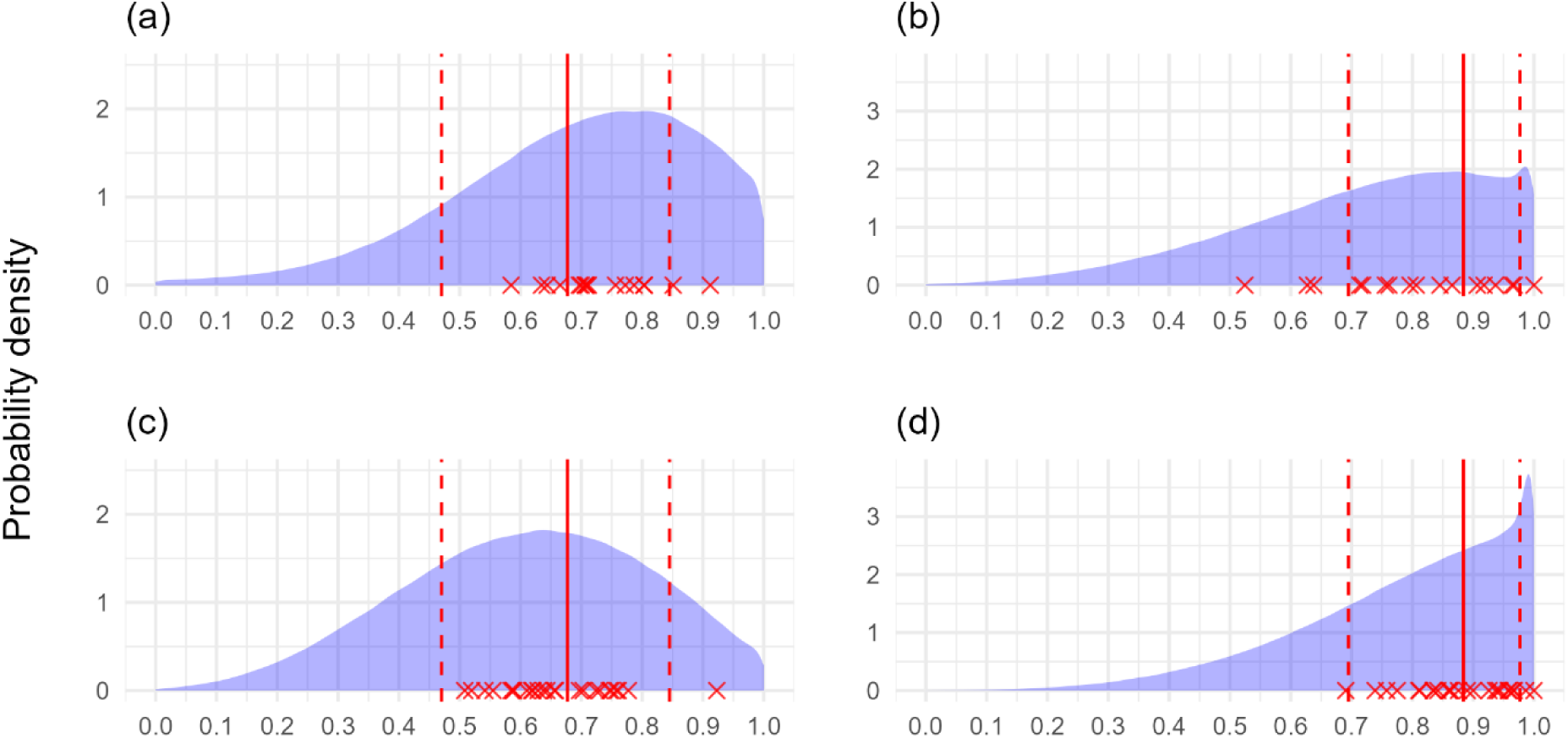
Workflow sensitivity and specificity distribution output using simulated data (80 trees, poor knowledge both symptoms). Probability density of estimated sensitivity and specificity of surveyors using poor sensitivity and specificity prior distributions with (a, b) no covariance model with no simulated covariance, (c, d) covariance model with simulated covariance. In both instances 80 trees each were assessed in higher and lower true disease prevalence locations by 25 surveyors, but 8/25 models did not adequately converge for the no covariance model and were discarded. Red crosses represent true surveyor sensitivity and specificity. Solid red line represents the 50^th^ percentile (median), dotted red lines represent the 5^th^ and 95^th^ percentiles of the distributions surveyor sensitivity and specificity were simulated from.

## Discussion

This study provides the first approach to derive quantitative estimates of the sensitivity and specificity of visual inspection for symptoms in plant health in the absence of a gold-standard reference dataset. This meets a key knowledge gap, since visual inspection is a cornerstone of plant health surveys, and lack of estimates for these parameters severely hinder adequate survey planning, interpretation of surveillance data, and instigation of effective outbreak management. Our findings highlight variation in ability between individual surveyors and different symptoms, thus emphasising the necessity of quantifying these parameters to optimise surveillance strategies in plant health. To facilitate this, our results demonstrate the utility of a workflow utilising Bayesian modelling to provide estimates of the sensitivity and specificity in the absence of rarely available gold standard data and informative prior information of these parameters.

Our results suggest fundamental surveillance metrics such as the number of visual assessments to confidently declare an area as pest free, understanding of the true pest prevalence given a positive detection, and reliability of positive detection [11] will vary depending on the visual symptom of the pest. Previous work highlights that the sensitivity of a detection method affects the optimal deployment of surveillance strategies for early detection of a pest or disease in the landscape, with lower sensitivity leading to surveillance focussed in higher-risk locations [36]. The variation in sensitivity in our results highlights that in plant health, not only the type of detection method (e.g. insect trapping vs visual detection), but the particular visual symptom on a host will influence the optimal survey locations for early detection.

In addition to informing AOD surveillance, in combination with epidemic spread models, the estimates of sensitivity presented in this study can be used to inform surveillance strategies for plant pests and diseases of major concern. For example, the emerald ash borer (EAB) beetle is another *Agrilus* species that has killed millions of ash trees in the US, and also presents with D-shaped exit holes [37] that are directly comparable to *A. biguttatus* exit holes associated with AOD. *Phytophthora* spp., are a major cause of forest disease globally, and frequently present with bleeding canker symptoms directly comparable to those examined in this study [38]. However, the sensitivity presented for the AOD symptoms represents the sensitivity of detecting symptoms. Therefore, for application to other pests and diseases both the asymptomatic period and frequency of asymptomatic hosts must be considered.

Our results also demonstrate that sensitivity of symptom detection is greater when hosts are more affected by a pest (i.e. greater symptom abundance). First, this means that the difficulty of early detection of pests in the landscape due to low abundance will be compounded by a lower sensitivity of detection of initial symptoms of a pest. Second, these results demonstrate that the pathology of a disease must be considered to optimally apply results.

Importantly, the surveyors in our study were citizen scientists, and not professional tree health surveyors. Citizen scientists received the same training prior to conducting AOD surveys, but still varied greatly in their ability to visually detect symptoms. Citizen science has an important role in plant health [14], yet it is not known how surveyors with different experience differ in their ability to detect plant pests and diseases. Previous work investigating the collection of phenological data, found that expert surveyors had less variability compared to citizen scientists, but citizen scientists (regardless of training) performed similar to the expert surveyors when recording the presence of abundant flowers and fruits [39]. Moreover, Pocock et al. [40] found that citizen scientist data agreed well with expert data when recording *Cameraria ohridella* leaf damage scores and counts of *C. ohridella* adult moths, but not when counting the rarer associated parasitoid wasp, which was attributed to a lack of familiarity with the rarer species. Together, in the context of plant health, these results suggest that frequent training would be beneficial for inexperienced surveyors, or for symptoms that are less commonly encountered. It has been shown that landowner reporting data and professional survey data provide similar estimates of AOD distribution data in the UK, and professional monitoring can be more resource efficient by using citizen science reports to guide surveys [41]. However, better understanding of the variation in surveyors’ ability, and the effect of training and experience are required for optimised survey design, and effective integration of different types of citizen science data with professional surveys.

The difficulty obtaining ‘gold-standard’ survey datasets prohibits the routine quantification of the sensitivity and specificity of visual inspection in plant health. Estimates of these parameters are possible through inoculation studies, but this is practically difficult to undertake for long-lived hosts (e.g. trees) and diseases that have multi-factorial causes or develop over a prolonged period (e.g. AOD). It is also challenging to ensure that the disease development from controlled inoculations is representative of disease development in the field, and that the presence of other ailments that can be mistaken for a pest or disease are appropriately represented under more controlled experimental conditions.

In our study, the expert survey data was assumed to be gold-standard because this expert has more than a decade of experience surveying these trees for AOD [20]. It is possible that the expert did not classify all trees correctly, although the high performance of some citizen scientists with comparatively limited experience (Fig 5), suggests the expert will have sensitivity and specificity of at least >0.90, which should have minimal impact on the findings. However, the sensitivity and specificity values from the AOD survey represent the probability of correctly classifying symptom presence or absence, rather than pest or disease presence or absence because the sensitivity and specificity parameters were calculated relative to expert visual detection of the symptoms.

The workflow outlined in this paper provides a practical solution to the problem of obtaining a gold-standard survey dataset. We show that with 25 surveyors, assessing a minimum of 80 plants each for two symptoms in lower (∼0.3) and higher (∼0.6) true disease prevalence locations, we can estimate the distribution of sensitivity and specificity for symptoms. It is also possible to estimate the sensitivity and specificity of individual surveyors, but it is not yet known how individual surveyors vary between surveys, and it would be practically and financially challenging to quantify each individual surveyor that will be visually inspecting plants. We therefore envisage a one-off organised survey event within the current range of a pest in higher and lower true prevalence sites that can be estimated from delimitation surveys following discovery of a pest [12]. The survey event will involve at least 25 surveyors that will record two prespecified symptoms of the pest of interest on each plant (at least 80 plants in each site). The minimum prior information required for subsequent modelling is knowledge of higher, or lower true disease prevalence sites, although greater prior knowledge of other parameters improves model accuracy. The output from the workflow will provide an estimate of the distribution of sensitivity and specificity for this group of surveyors, to then be applied to different groups of surveyors with a similar level of experience. It is recommended surveyors have similar experience given that distributions of sensitivity and specificity will likely vary with experience. These estimates of parameter distributions can then be used to determine fundamental surveillance metrics [11] described earlier, as well as for optimisation of survey locations to maximise the likelihood of early detection [36].

The sensitivity output from the workflow represents the probability of correctly identifying pest or disease presence from visual inspection for a given symptom within the surveyed areas. Post-hoc analyses utilising knowledge of the pathology and epidemiology of a disease can therefore be used to increase the accuracy of sensitivity estimates for application to other areas by accounting for spatial and temporal variation in both the expected proportions of asymptomatic hosts and symptom severity. Unreliable model estimates were obtained in our analyses when sensitivity and specificity values were at the tail of prior distributions, but this can be addressed by appropriate specification of prior information in Bayesian analyses using a combination of ‘common biological sense’ and sensitivity analyses of priors [42].

Previous work demonstrates that models which don’t account for non-independence of sensitivity and specificity between tests by failing to include covariance parameters in the model perform poorly [29]. This non-independence is likely to be present when tests examine comparable biological aspects of a disease [29]. For example, the sensitivity and specificity of detection of a pest from a necrotic lesion via an ELISA test for a pest antigen and via qPCR for pest DNA are unlikely to be independent. For visual detection of pests, this is particularly likely to occur when symptoms co-occur spatially and temporally. However, our results highlight that including additional covariance parameters in the model incurs negligible penalty in the example used here. This likely reflects the benefit of informative prior knowledge of the true disease prevalence on sites, the fact that the true disease prevalence was very different between sites, and that a large number of hosts were surveyed on each site (at least 80 trees). Given the workflow presented in this paper can provide sensitivity and specificity estimates with the presence of covariance in the data and model, and a potentially large penalty is incurred in models which falsely exclude covariance [29], we recommend including a covariance parameter in the model unless convincing data suggests otherwise. In future, more informative prior information on the covariance parameters for specific systems would strengthen model performance.

### Conclusions

This study demonstrates that the sensitivity and specificity of visual inspection varies between plant health symptoms and between surveyors. The data generated in our study for the inspection of AOD symptoms can be applied to surveillance for pests which present similar symptoms, such as EAB and *Phytophthora* spp. However, given the ability to detect symptoms improves with symptom frequency, further work investigating the temporal pattern of symptom appearance (including asymptomatic period) would be highly valuable for plant health surveillance. Future work should also investigate the large surveyor variation, and understand the effect of training and experience of surveyors to allow better integration of citizen science activities with professional surveys. The variation in sensitivity and specificity between symptoms and surveyors means that: (1) the reliability of survey reports will vary between pests and surveyors; (2) different numbers of survey records will be required to declare different pests absent with the same confidence; (3) the true incidence given a positive detection will depend on the surveyor and symptom of the pest, not just the epidemiology; (4) survey locations to maximise the likelihood of early detection will differ depending on the visual symptom of a pest. To apply this knowledge, the workflow presented in this paper enables quantification of the sensitivity and specificity for visual plant health surveys in the absence of rarely available ‘gold-standard’ datasets. This addresses a major knowledge gap, which will enable the calculation of fundamental surveillance metrics, and improve risk-based surveillance strategies in plant health. Further adaptation of these methods can be used for a broad range of visual environmental surveillance activities.

## Acknowledgements

We would like to thank all involved in the AOD survey days.

## Code and data availability

The code and data used for this paper are available from: https://github.com/MCombess/Plant_Health_sens_spec_workflow and are archived: DOI: 10.5281/zenodo.14975201

## Author contributions

**Conceptualization:** Matt Combes, Stephen Parnell, Nathan Brown, Alexander Mastin.

**Formal analysis:** Matt Combes, Stephen Parnell, Robin N. Thompson, Alexander Mastin.

**Funding acquisition:** Stephen Parnell, Nathan Brown, Peter Crow.

**Investigation:** Nathan Brown, Peter Crow.

**Methodology:** Matt Combes, Stephen Parnell, Nathan Brown, Robin N. Thompson, Alexander Mastin.

**Project administration:** Stephen Parnell, Nathan Brown.

**Supervision:** Stephen Parnell

**Visualization:** Matt Combes

**Writing – original draft:** Matt Combes, Stephen Parnell.

**Writing – review & editing:** Matt Combes, Stephen Parnell, Nathan Brown, Robin N. Thompson, Alexander Mastin, Peter Crow.

## Supporting information

Additional supporting information can be found online in the Supporting Information section at the end of this article.

**S1 File. Details on pre-survey AOD training and citizen scientists’ tree and tree health experience scored from a questionnaire.**

**S2 File. Citizen scientist surveyor sensitivity and specificity values.**

**S3 File. Output statistics from model runs (individuals and distributions) with mean squared error values, and details of the prior probability distributions.**

**S1 Fig.**
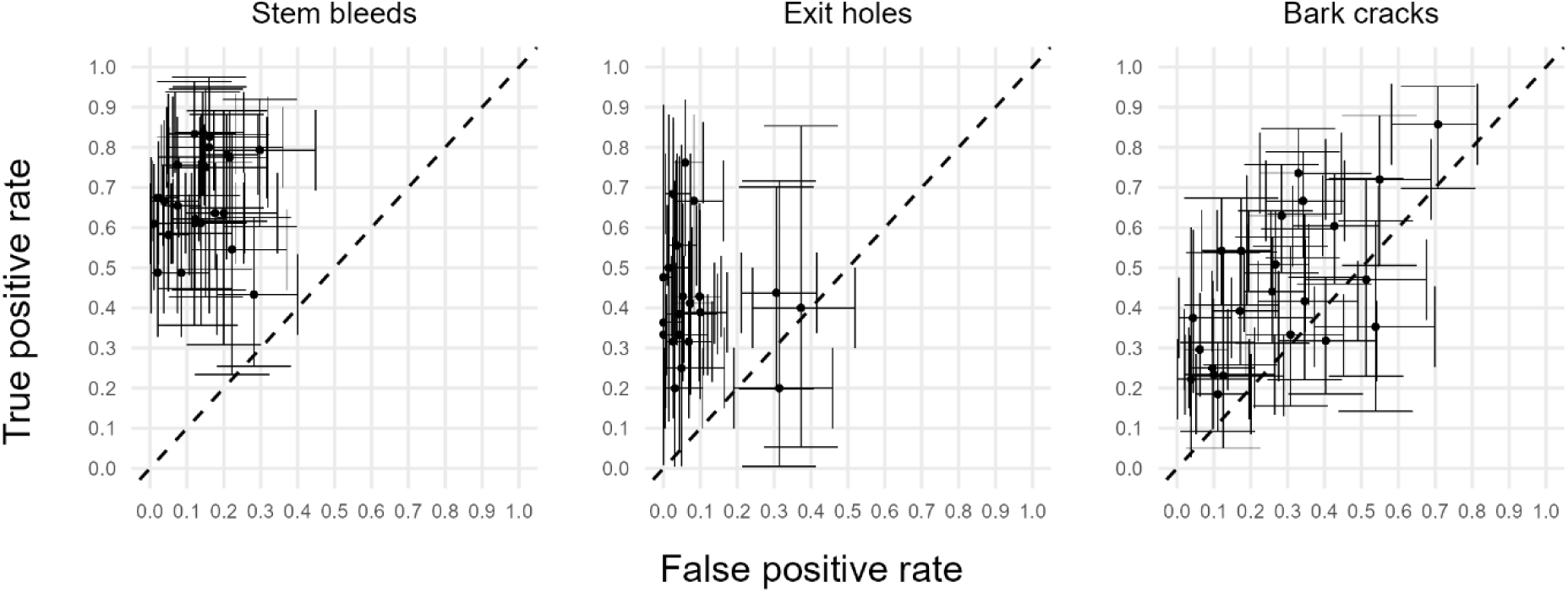
Receiver operating characteristic plots for three externally visible AOD symptoms. The presence/ absence of bleeds/ weeping patches on the main stem, *Agrilus* beetle exit holes and cracks between bark plates were assessed by 23 surveyors on up to 175 oak trees. True positive rate and false positive rate were calculated using expert data as a reference standard. Error bars represent 95% confidence intervals.

**S2 Fig.**
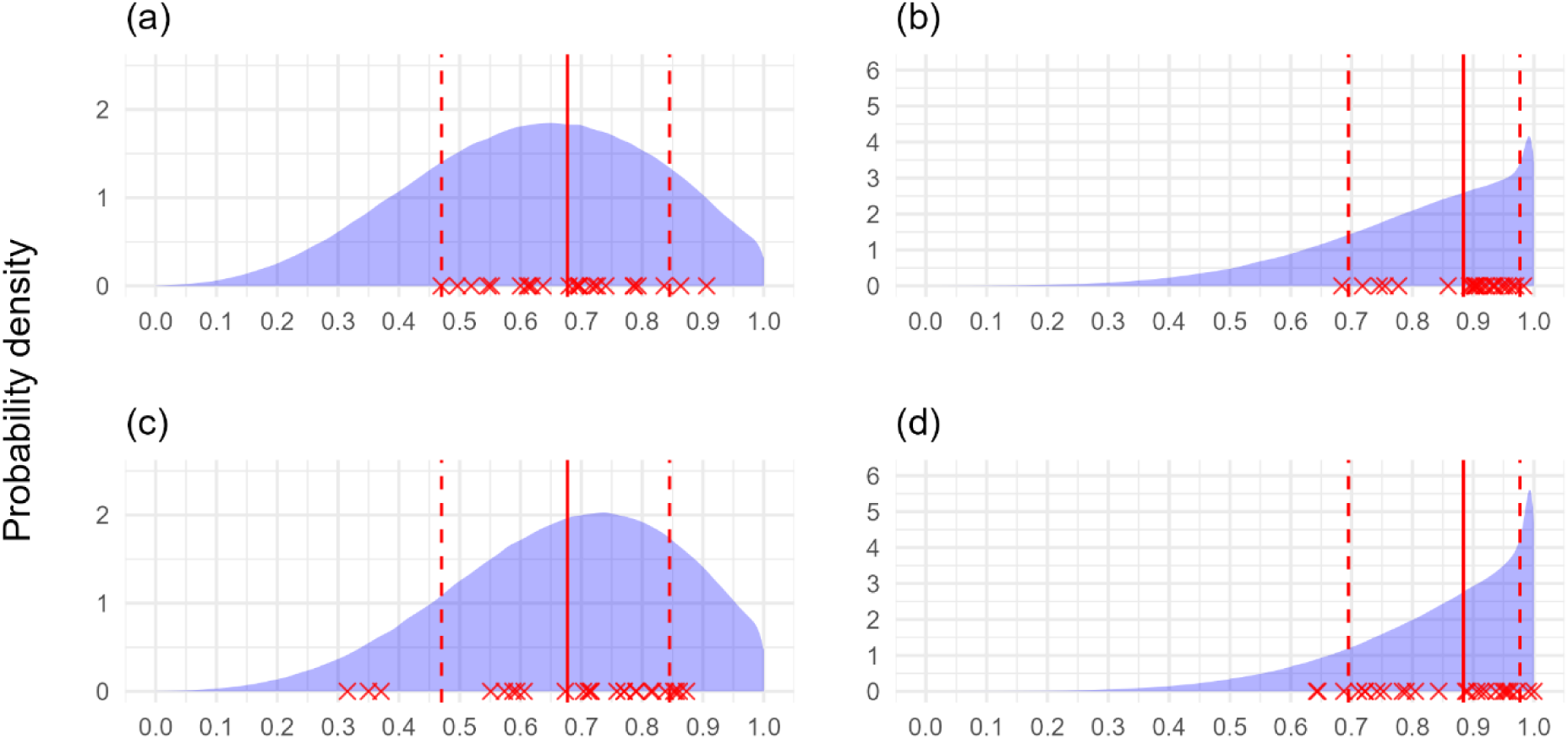
Workflow sensitivity and specificity distribution output using simulated data (80 trees, very good knowledge one symptom). Probability density of estimated sensitivity and specificity of surveyors using poor sensitivity and specificity prior distributions for symptom one, and very good sensitivity and specificity prior distributions for symptom two. Workflow output is presented for symptom one with (a, b) no covariance model with no simulated covariance, (c, d) covariance model with simulated covariance. In both instances 80 trees each were assessed in higher and lower true disease prevalence locations by 25 surveyors. Red crosses represent true surveyor sensitivity and specificity. Solid red line represents the 50^th^ percentile (median), dotted red lines represent the 5^th^ and 95^th^ percentiles of the distributions surveyor sensitivity and specificity were simulated from.

**S3 Fig.**
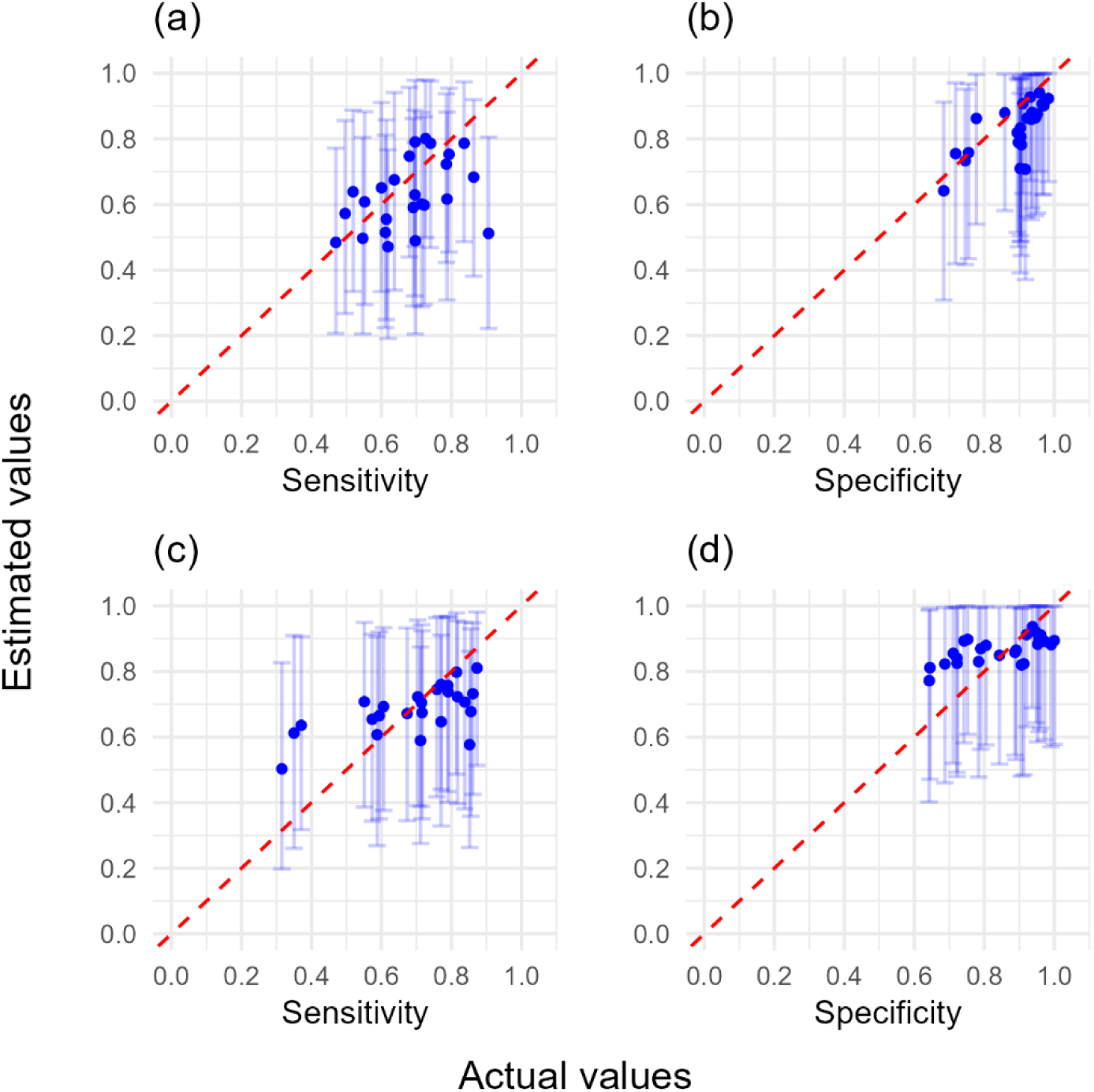
The workflow estimated sensitivity and specificity for simulated surveyors against actual sensitivity and specificity values (80 trees, very good knowledge one symptom). The workflow used poor sensitivity and specificity prior distributions for symptom one, and very good sensitivity and specificity prior distributions for symptom two. Workflow output is presented for symptom one with (a, b) no covariance model with no simulated covariance, (c, d) covariance model with simulated covariance. In both instances 80 trees each were assessed in higher and lower true disease prevalence locations. Error bars represent 95% confidence intervals, and dashed line represents perfect agreement between workflow estimated and actual values.

**S4 Fig.**
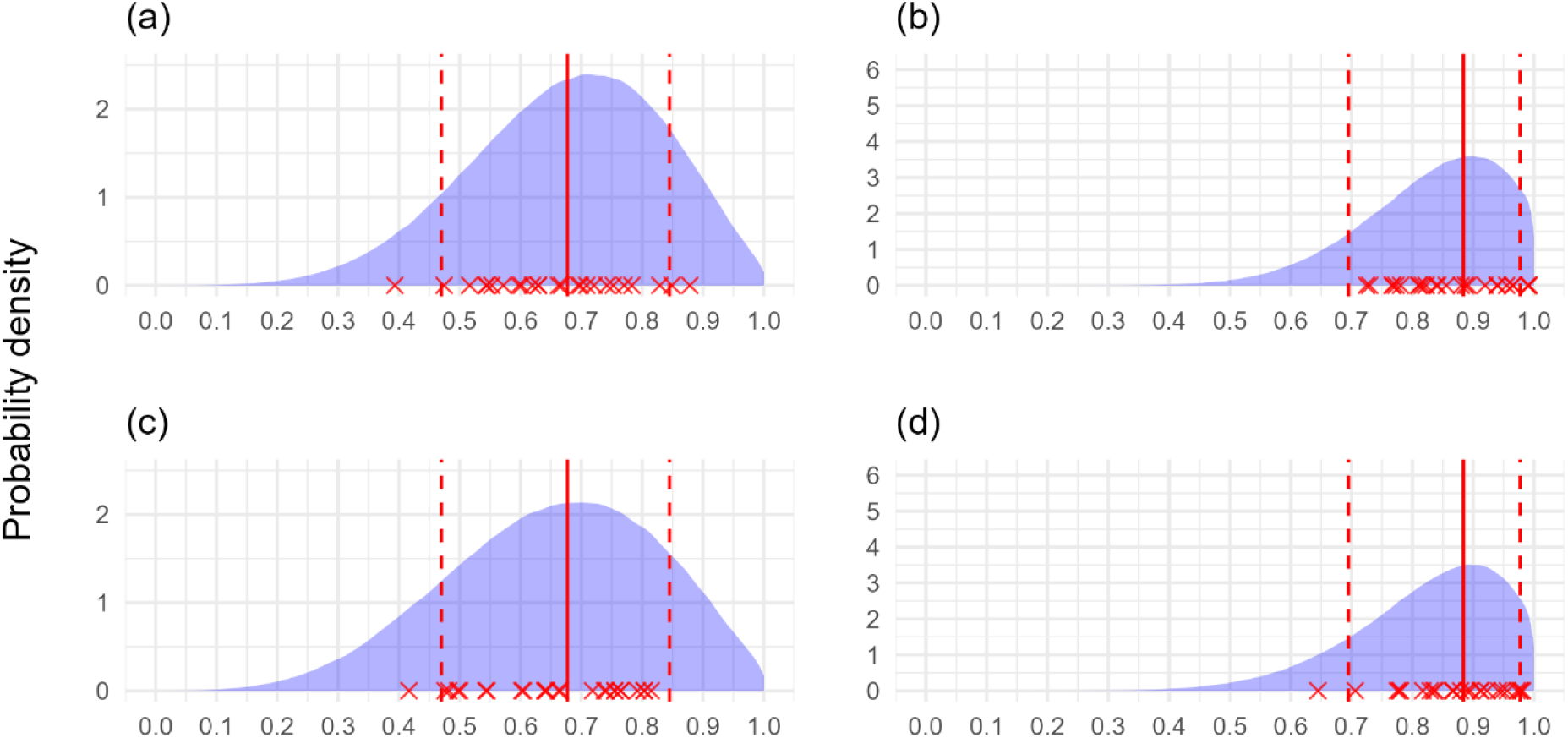
Workflow sensitivity and specificity distribution output using simulated data (80 trees, good knowledge both symptoms). Probability density of estimated sensitivity and specificity of surveyors using good sensitivity and specificity prior distributions with (a, b) no covariance model with no simulated covariance, (c, d) covariance model with simulated covariance. In both instances 80 trees each were assessed in higher and lower true disease prevalence locations by 25 surveyors. Red crosses represent true surveyor sensitivity and specificity. Solid red line represents the 50^th^ percentile (median), dotted red lines represent the 5^th^ and 95^th^ percentiles of the distributions surveyor sensitivity and specificity were simulated from.

**S5 Fig.**
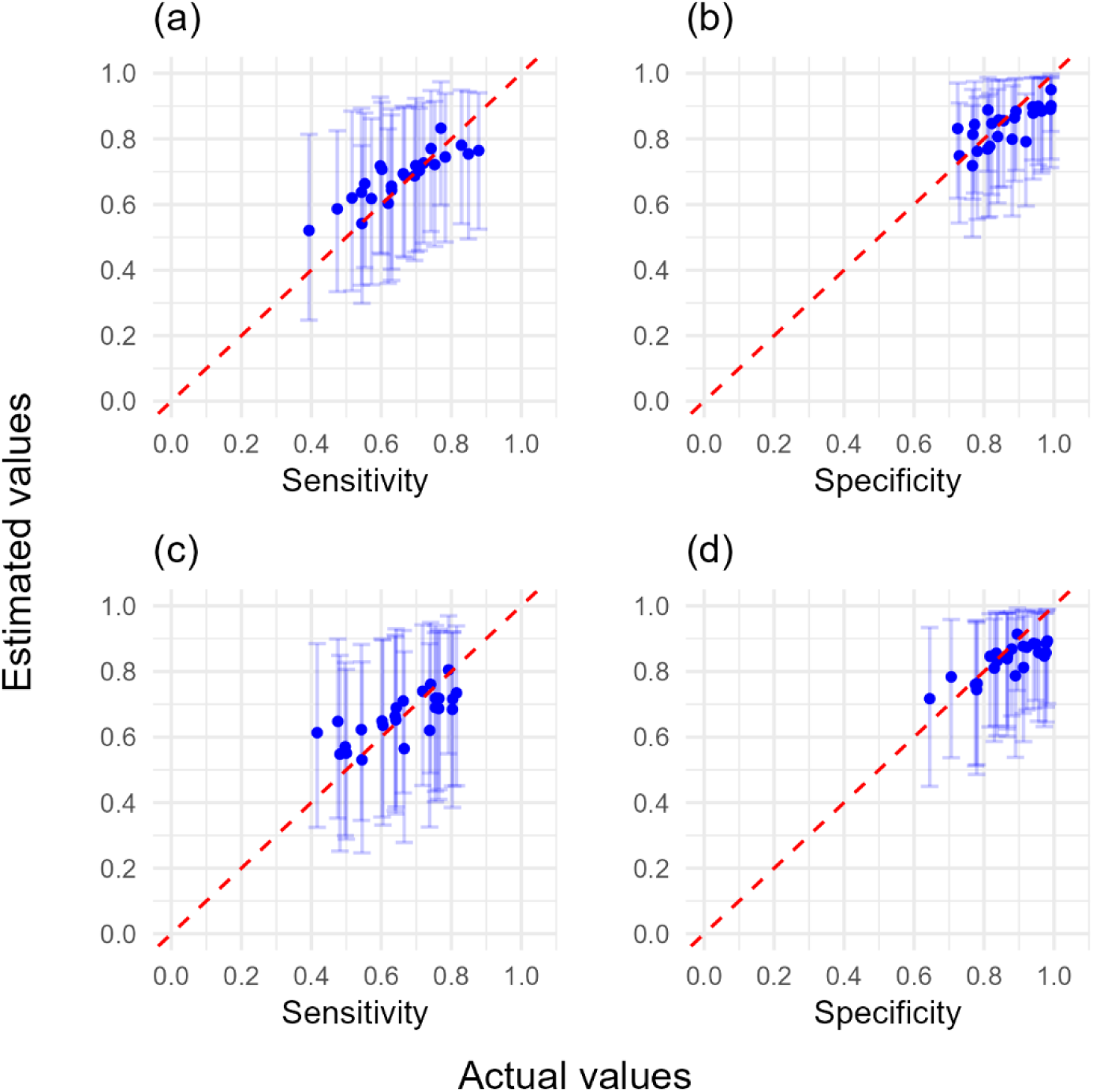
The workflow estimated sensitivity and specificity for simulated surveyors against actual sensitivity and specificity values (80 trees, good knowledge both symptoms). The workflow used good sensitivity and specificity prior distributions with (a, b) no covariance model with no simulated covariance, (c, d) covariance model with simulated covariance. In both instances 80 trees each were assessed in higher and lower true disease prevalence locations. Error bars represent 95% confidence intervals, and dashed line represents perfect agreement between workflow estimated and actual values.

**S6 Fig.**
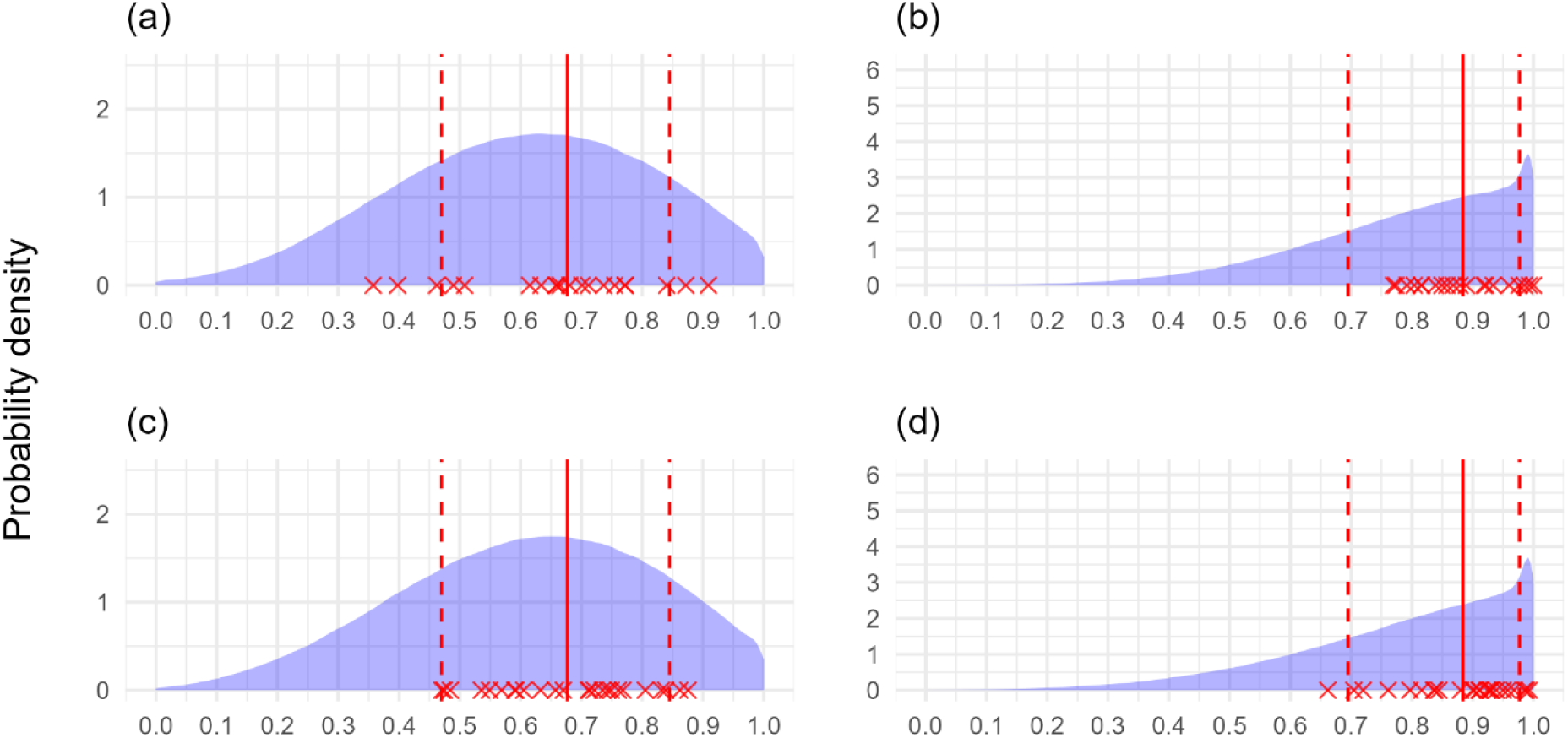
Workflow sensitivity and specificity distribution output using simulated data (100 trees, poor knowledge both symptoms). Probability density of estimated sensitivity and specificity of surveyors using poor sensitivity and specificity prior distributions with (a, b) no covariance model with no simulated covariance, (c, d) covariance model with simulated covariance. In both instances 100 trees each were assessed in higher and lower true disease prevalence locations by 25 surveyors, but 5/25 models did not adequately converge for the no covariance model and were discarded. Red crosses represent true surveyor sensitivity and specificity. Solid red line represents the 50^th^ percentile (median), dotted red lines represent the 5^th^ and 95^th^ percentiles of the distributions surveyor sensitivity and specificity were simulated from.

**S7 Fig.**
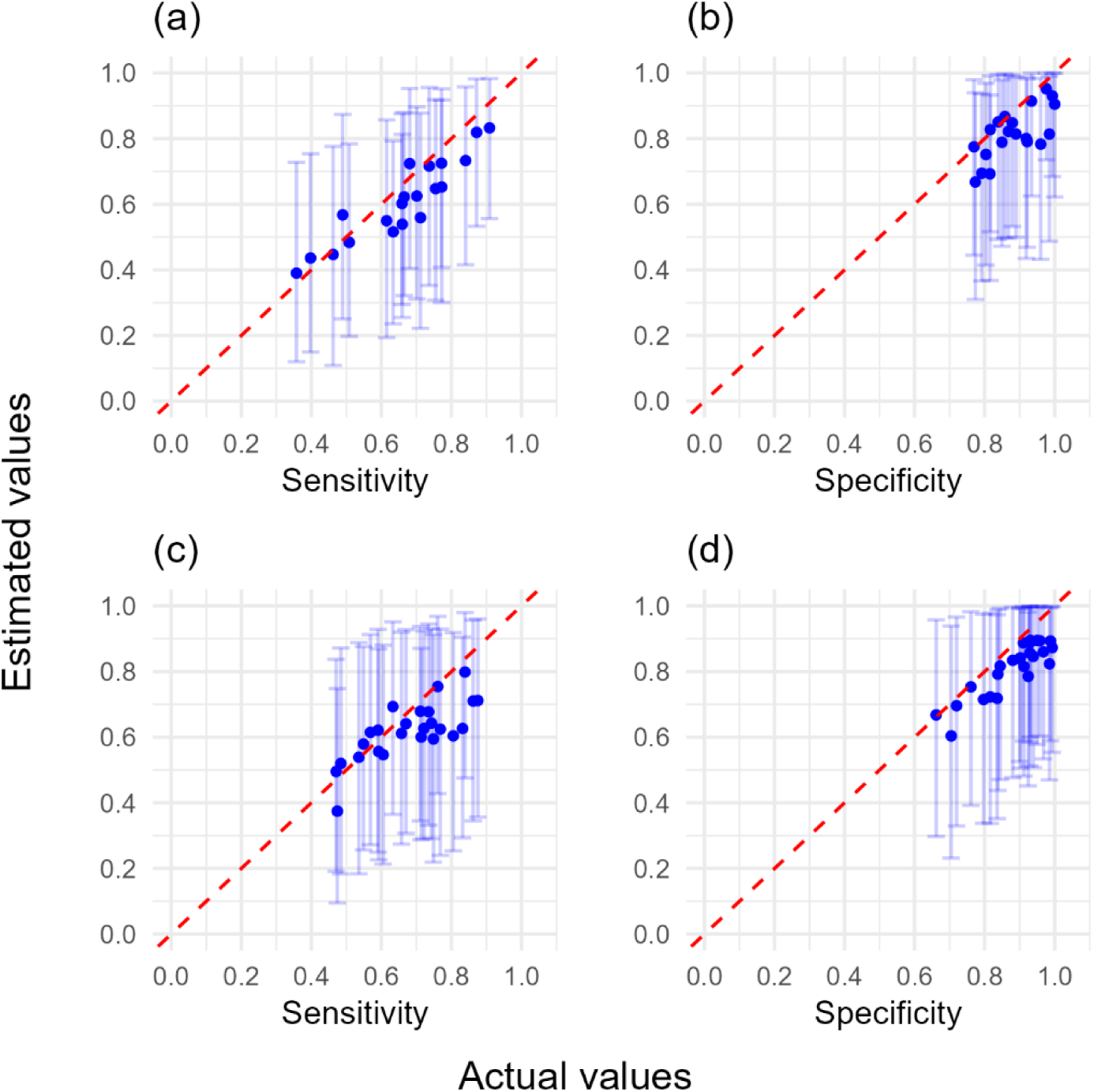
The workflow estimated sensitivity and specificity for simulated surveyors against actual sensitivity and specificity values (100 trees, poor knowledge both symptoms). The workflow used poor sensitivity and specificity prior distributions with (a, b) no covariance model with no simulated covariance, (c, d) covariance model with simulated covariance. In both instances 100 trees each were assessed in higher and lower true disease prevalence locations. Error bars represent 95% confidence intervals, and dashed line represents perfect agreement between workflow estimated and actual values.

**S8 Fig.**
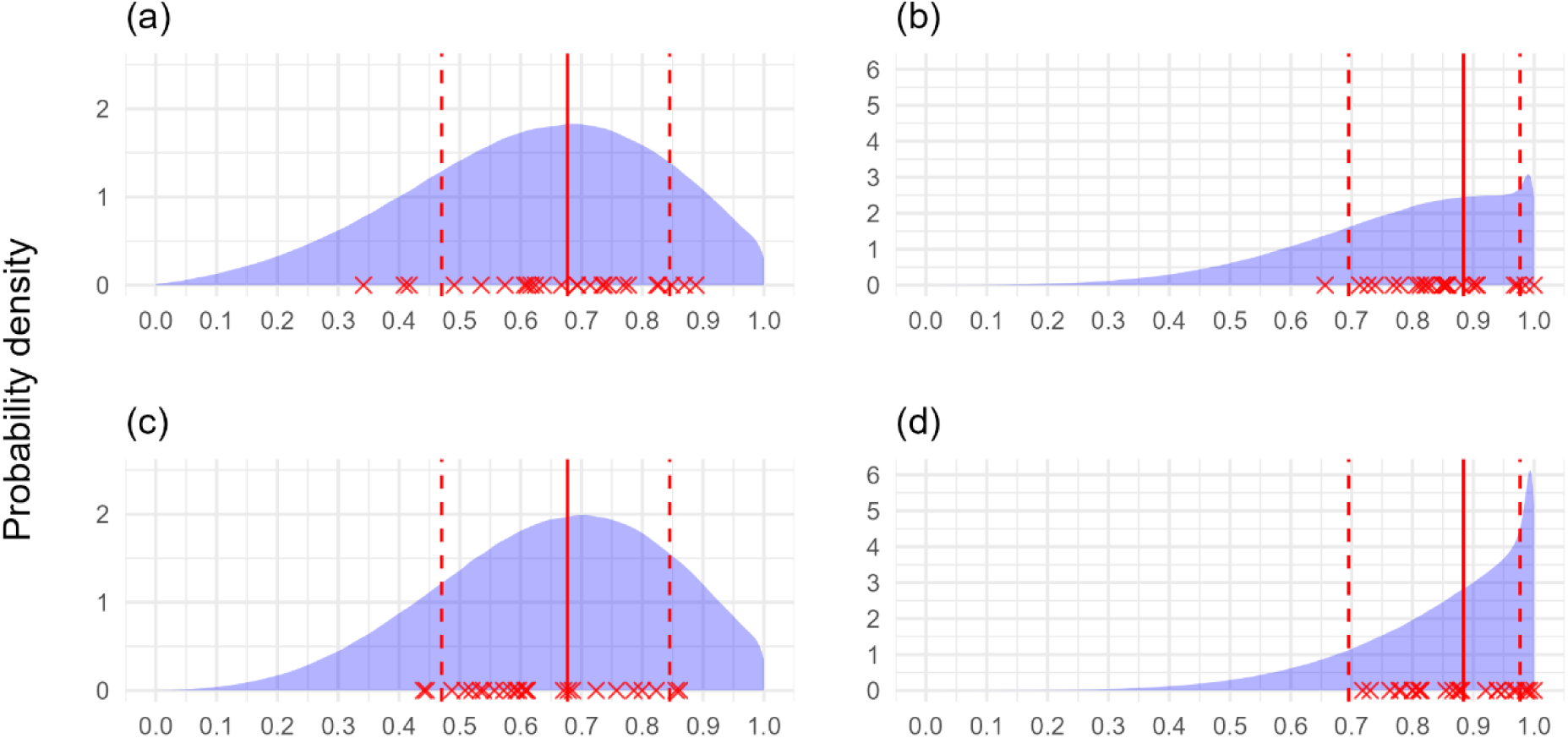
Workflow sensitivity and specificity distribution output using simulated data (100 trees, very good knowledge one symptom). Probability density of estimated sensitivity and specificity of surveyors using poor sensitivity and specificity prior distributions for symptom one, and very good sensitivity and specificity prior distributions for symptom two. Workflow output is presented for symptom one with (a, b) no covariance model with no simulated covariance, (c, d) covariance model with simulated covariance. In both instances 100 trees each were assessed in higher and lower true disease prevalence locations by 25 surveyors. Red crosses represent true surveyor sensitivity and specificity. Solid red line represents the 50^th^ percentile (median), dotted red lines represent the 5^th^ and 95^th^ percentiles of the distributions surveyor sensitivity and specificity were simulated from.

**S9 Fig.**
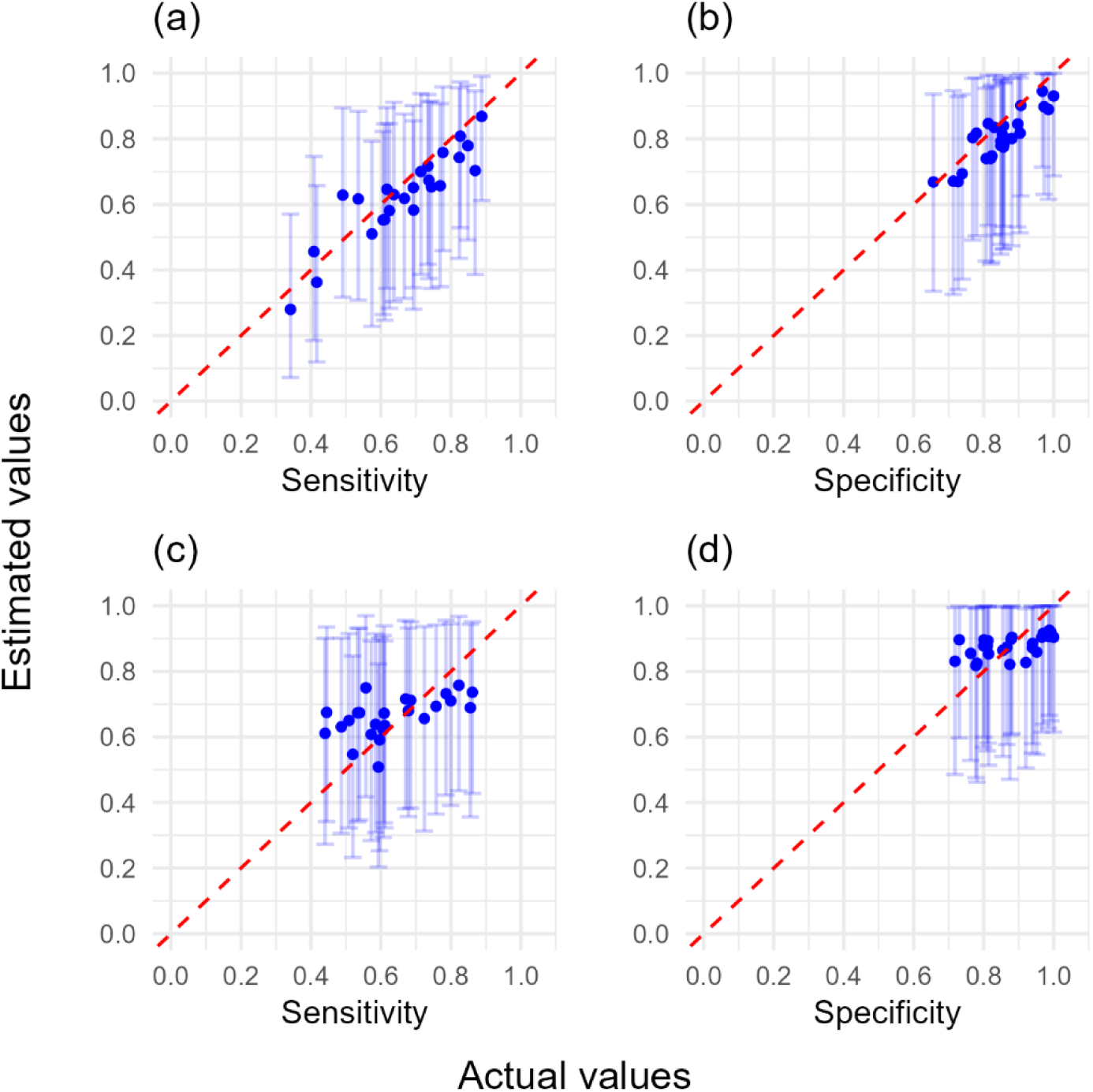
The workflow estimated sensitivity and specificity for simulated surveyors against actual sensitivity and specificity values (100 trees, very good knowledge one symptom). The workflow used poor sensitivity and specificity prior distributions for symptom one, and very good sensitivity and specificity prior distributions for symptom two. Workflow output is presented for symptom one with (a, b) no covariance model with no simulated covariance, (c, d) covariance model with simulated covariance. In both instances 100 trees each were assessed in higher and lower true disease prevalence locations. Error bars represent 95% confidence intervals, and dashed line represents perfect agreement between workflow estimated and actual values.

**S10 Fig.**
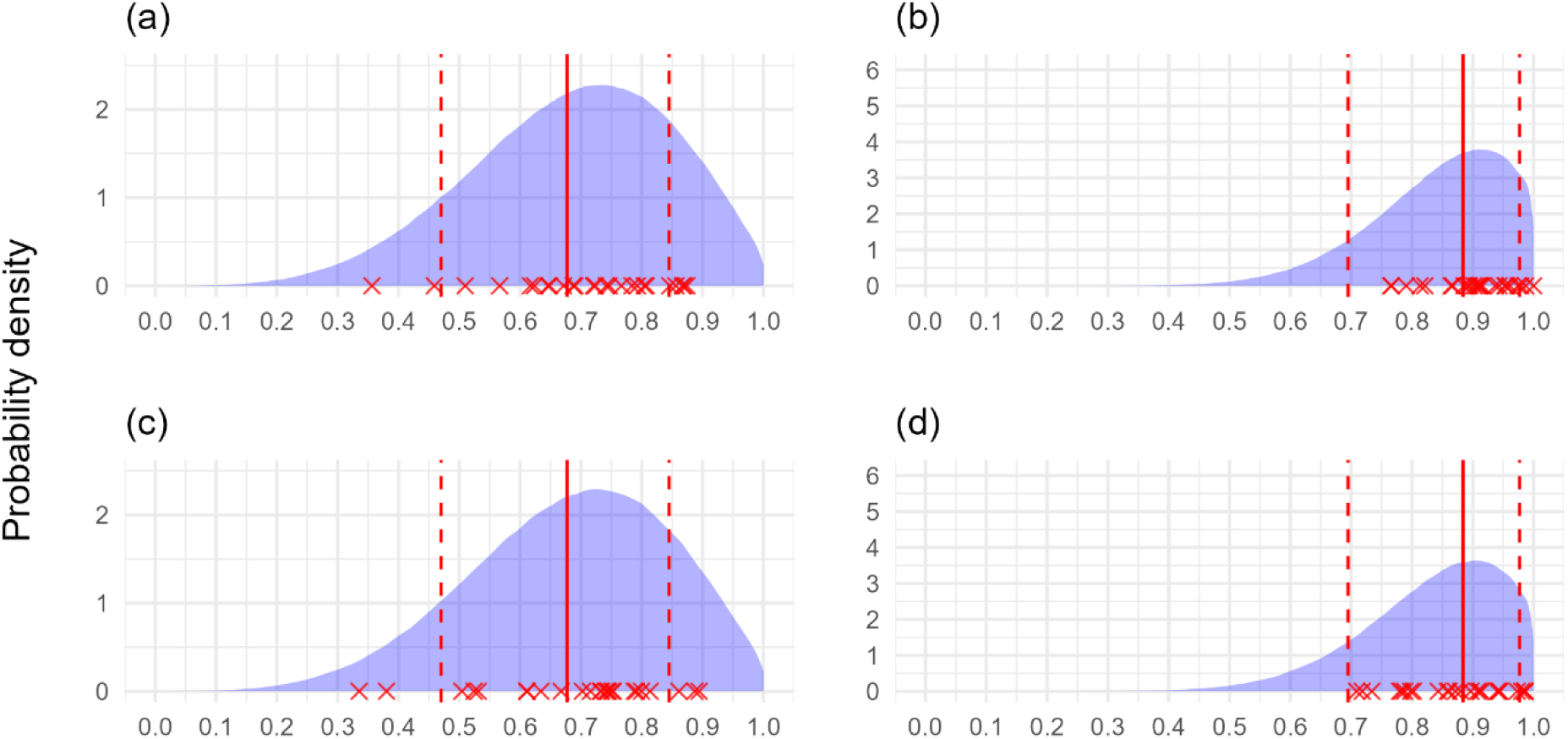
Workflow sensitivity and specificity distribution output using simulated data (100 trees, good knowledge both symptoms). Probability density of estimated sensitivity and specificity of surveyors using good sensitivity and specificity prior distributions with (a, b) no covariance model with no simulated covariance, (c, d) covariance model with simulated covariance. In both instances 100 trees each were assessed in higher and lower true disease prevalence locations by 25 surveyors. Red crosses represent true surveyor sensitivity and specificity. Solid red line represents the 50^th^ percentile (median), dotted red lines represent the 5^th^ and 95^th^ percentiles of the distributions surveyor sensitivity and specificity were simulated from.

**S11 Fig.**
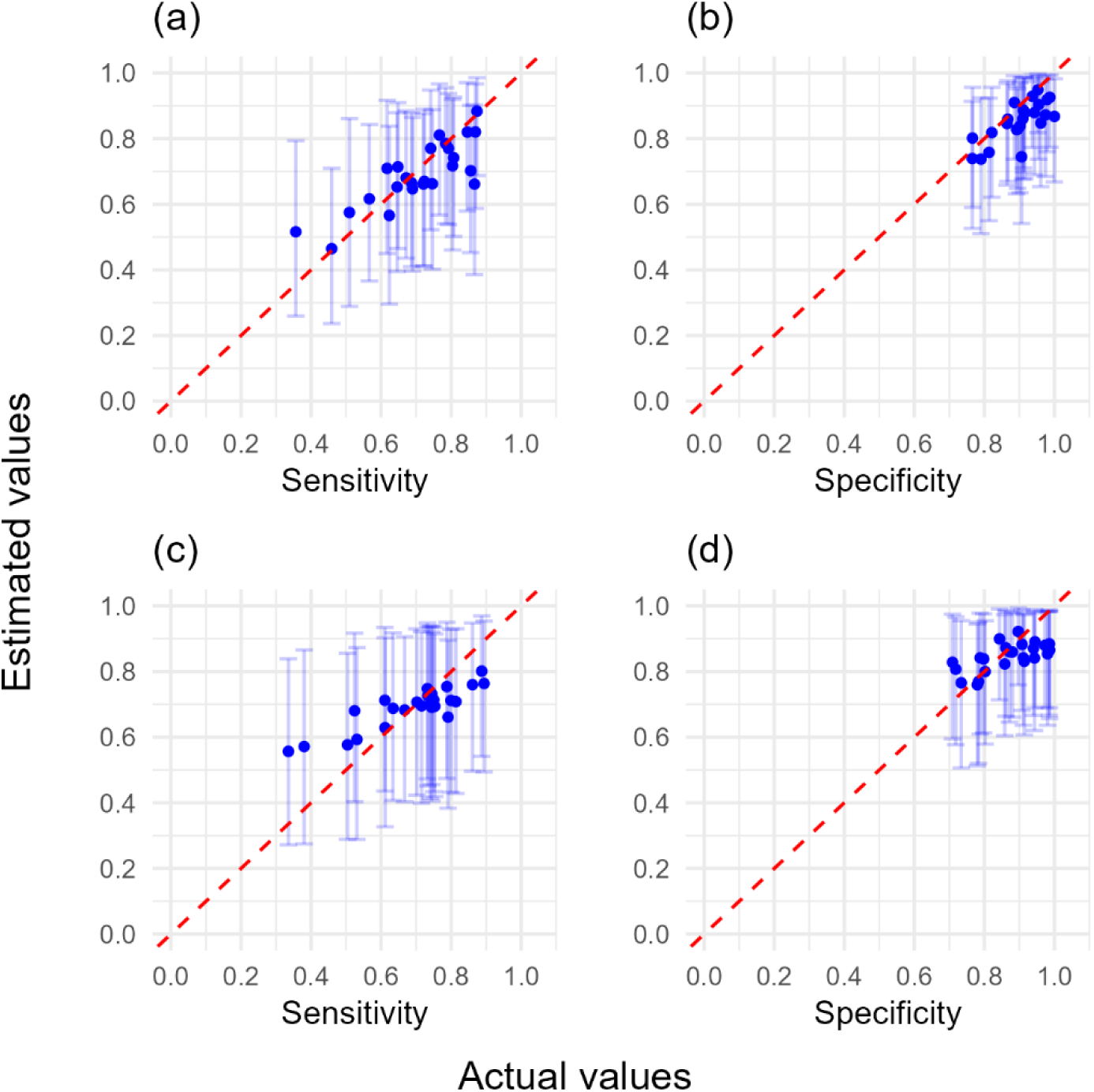
The workflow estimated sensitivity and specificity for simulated surveyors against actual sensitivity and specificity values (100 trees, good knowledge both symptoms). The workflow used good sensitivity and specificity prior distributions with (a, b) no covariance model with no simulated covariance, (c, d) covariance model with simulated covariance. In both instances 100 trees each were assessed in higher and lower true disease prevalence locations. Error bars represent 95% confidence intervals, and dashed line represents perfect agreement between workflow estimated and actual values.

**S12 Fig.**
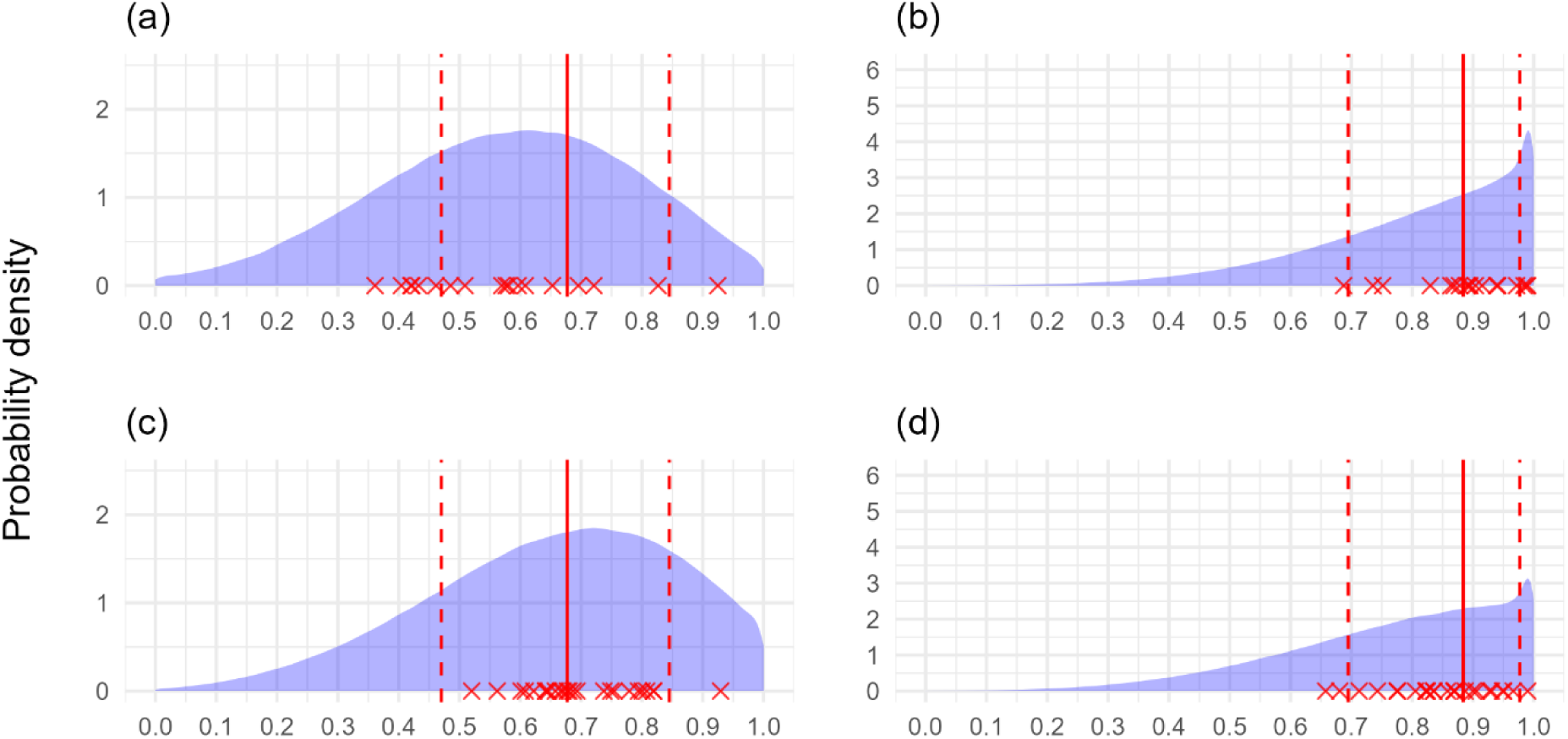
Workflow sensitivity and specificity distribution output using simulated data (120 trees, poor knowledge both symptoms). Probability density of estimated sensitivity and specificity of surveyors using poor sensitivity and specificity prior distributions with (a, b) no covariance model with no simulated covariance, (c, d) covariance model with simulated covariance. In both instances 120 trees each were assessed in higher and lower true disease prevalence locations by 25 surveyors, but 7/25 models did not adequately converge for the no covariance model and were discarded. Red crosses represent true surveyor sensitivity and specificity. Solid red line represents the 50^th^ percentile (median), dotted red lines represent the 5^th^ and 95^th^ percentiles of the distributions surveyor sensitivity and specificity were simulated from.

**S13 Fig.**
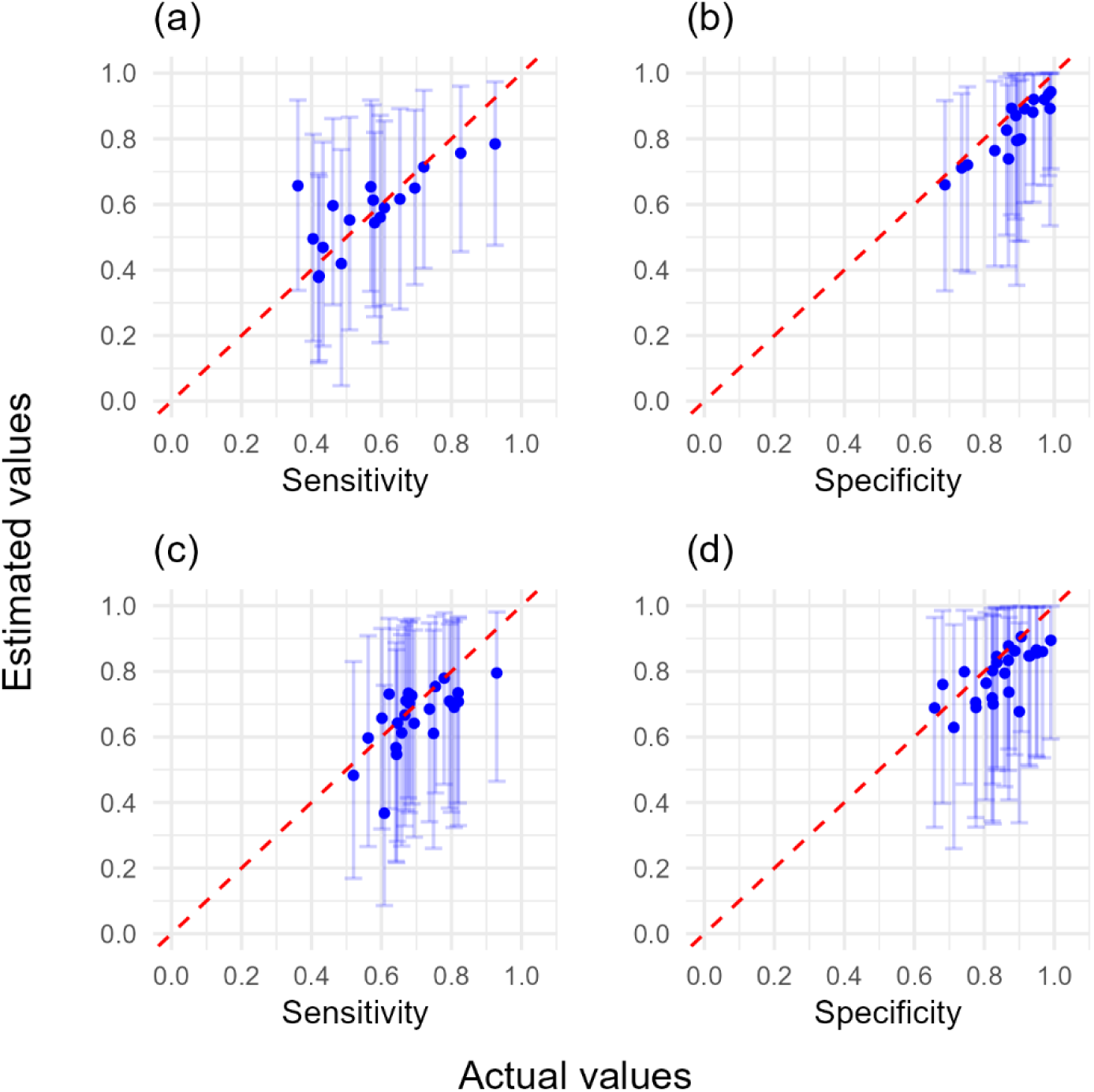
The workflow estimated sensitivity and specificity for simulated surveyors against actual sensitivity and specificity values (120 trees, poor knowledge both symptoms). The workflow used poor sensitivity and specificity prior distributions with (a, b) no covariance model with no simulated covariance, (c, d) covariance model with simulated covariance. In both instances 120 trees each were assessed in higher and lower true disease prevalence locations. Error bars represent 95% confidence intervals, and dashed line represents perfect agreement between workflow estimated and actual values.

**S14 Fig.**
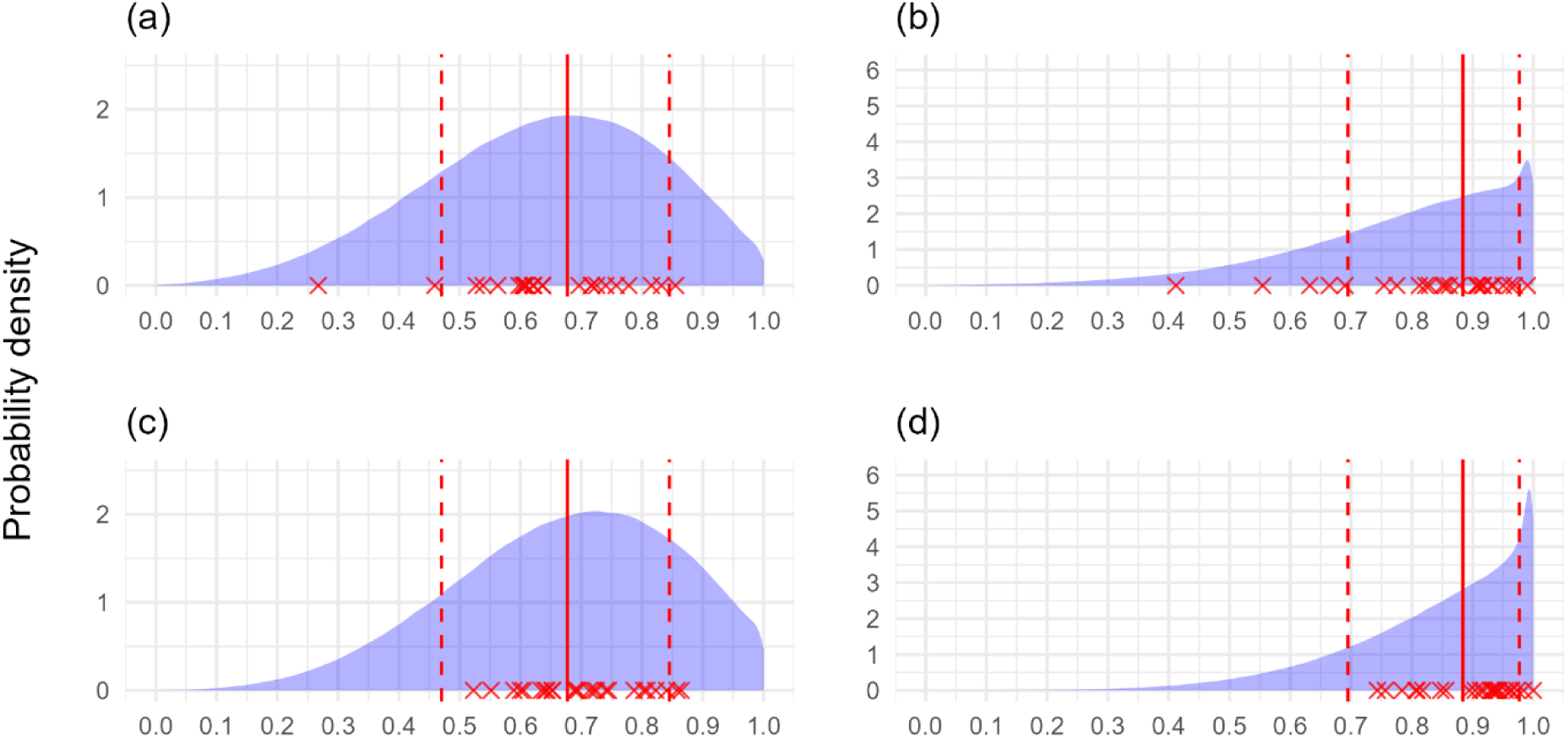
Workflow sensitivity and specificity distribution output using simulated data (120 trees, very good knowledge one symptom). Probability density of estimated sensitivity and specificity of surveyors using poor sensitivity and specificity prior distributions for symptom one, and very good sensitivity and specificity prior distributions for symptom two. Workflow output is presented for symptom one with (a, b) no covariance model with no simulated covariance, (c, d) covariance model with simulated covariance. In both instances 120 trees each were assessed in higher and lower true disease prevalence locations by 25 surveyors. Red crosses represent true surveyor sensitivity and specificity. Solid red line represents the 50^th^ percentile (median), dotted red lines represent the 5^th^ and 95^th^ percentiles of the distributions surveyor sensitivity and specificity were simulated from.

**S15 Fig.**
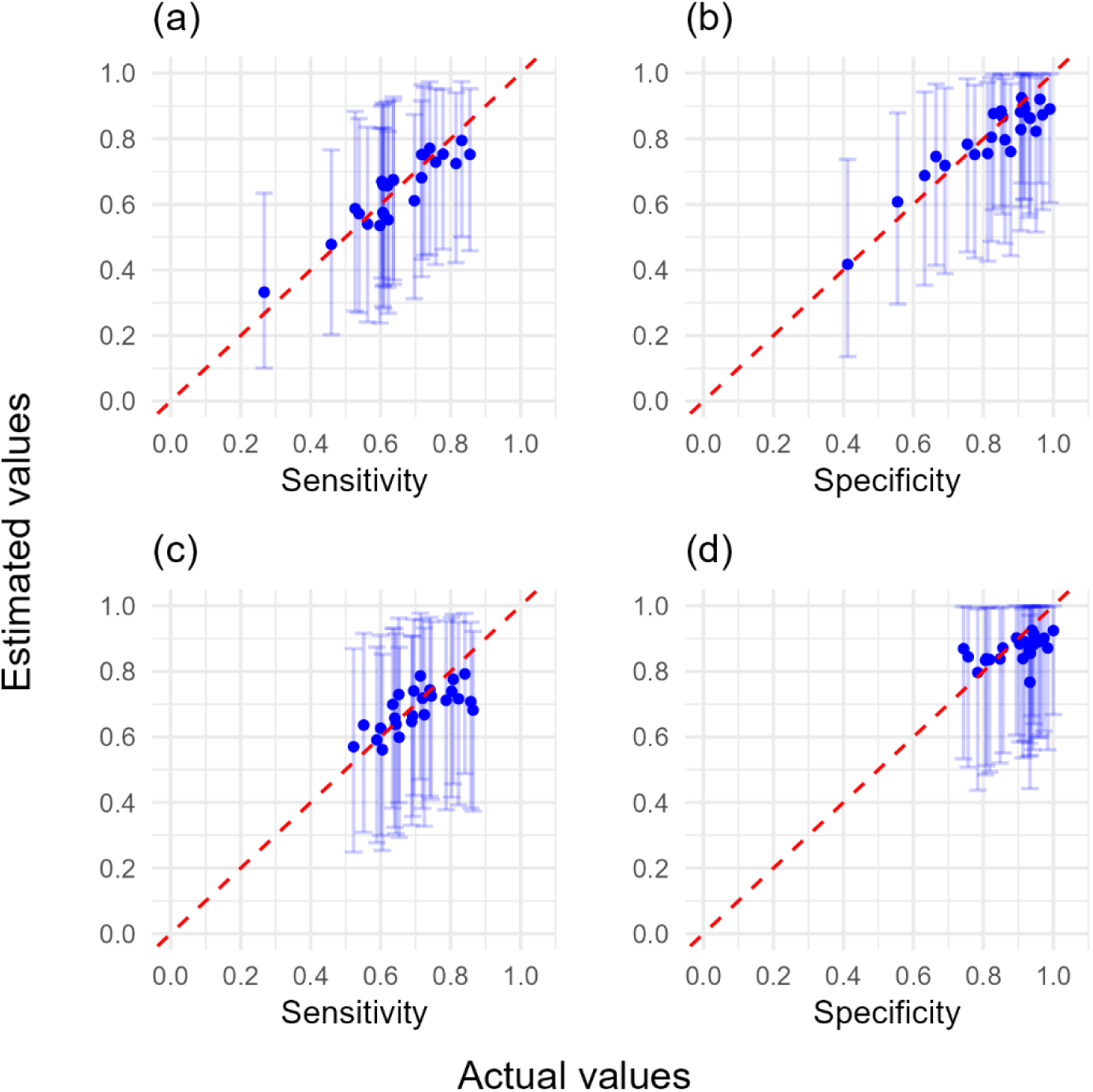
The workflow estimated sensitivity and specificity for simulated surveyors against actual sensitivity and specificity values (120 trees, very good knowledge one symptom). The workflow used poor sensitivity and specificity prior distributions for symptom one, and very good sensitivity and specificity prior distributions for symptom two. Workflow output is presented for symptom one with (a, b) no covariance model with no simulated covariance, (c, d) covariance model with simulated covariance. In both instances 120 trees each were assessed in higher and lower true disease prevalence locations. Error bars represent 95% confidence intervals, and dashed line represents perfect agreement between workflow estimated and actual values.

**S16 Fig.**
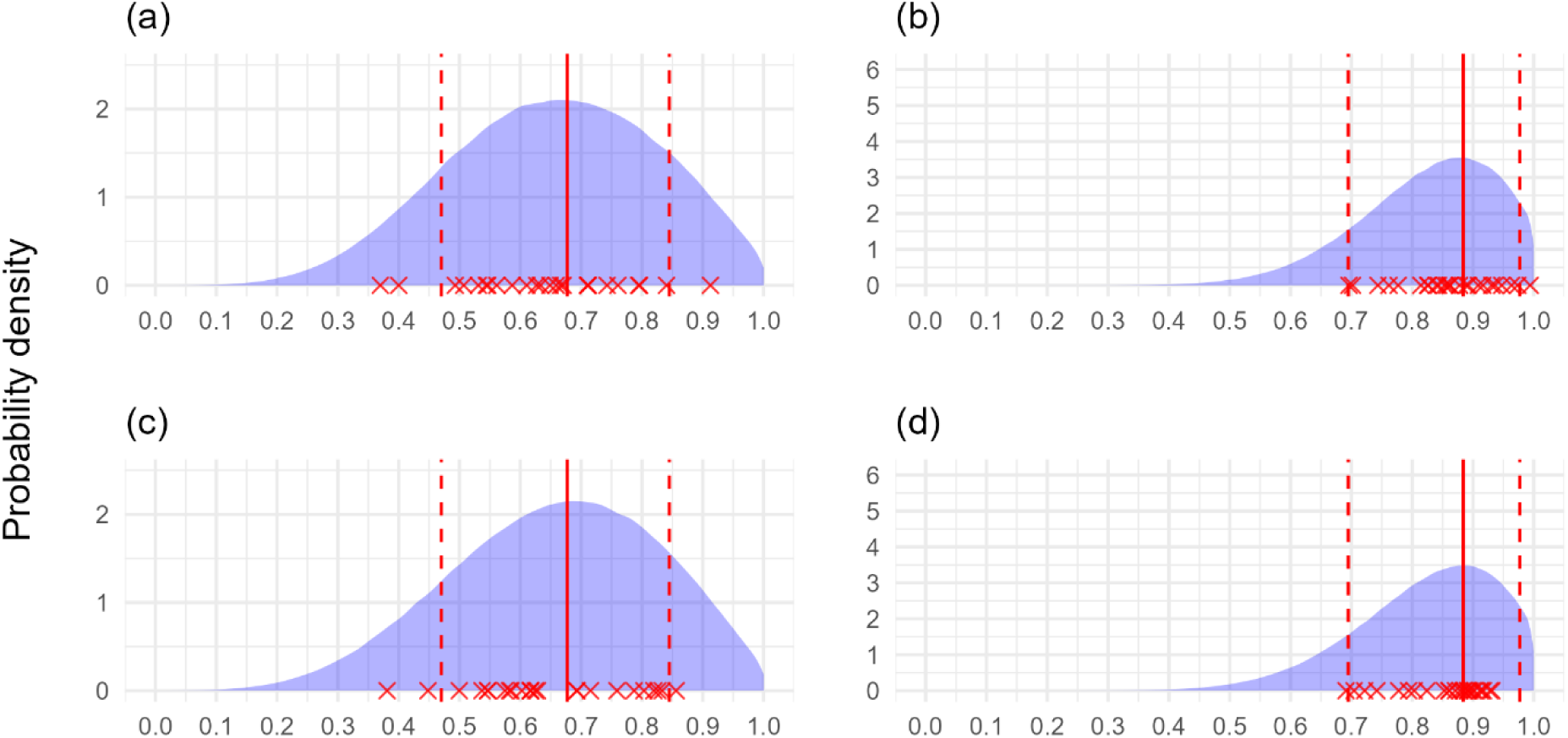
Workflow sensitivity and specificity distribution output using simulated data (120 trees, good knowledge both symptoms). Probability density of estimated sensitivity and specificity of surveyors using good sensitivity and specificity prior distributions with (a, b) no covariance model with no simulated covariance, (c, d) covariance model with simulated covariance. In both instances 120 trees each were assessed in higher and lower true disease prevalence locations by 25 surveyors. Red crosses represent true surveyor sensitivity and specificity. Solid red line represents the 50^th^ percentile (median), dotted red lines represent the 5^th^ and 95^th^ percentiles of the distributions surveyor sensitivity and specificity were simulated from.

**S17 Fig.**
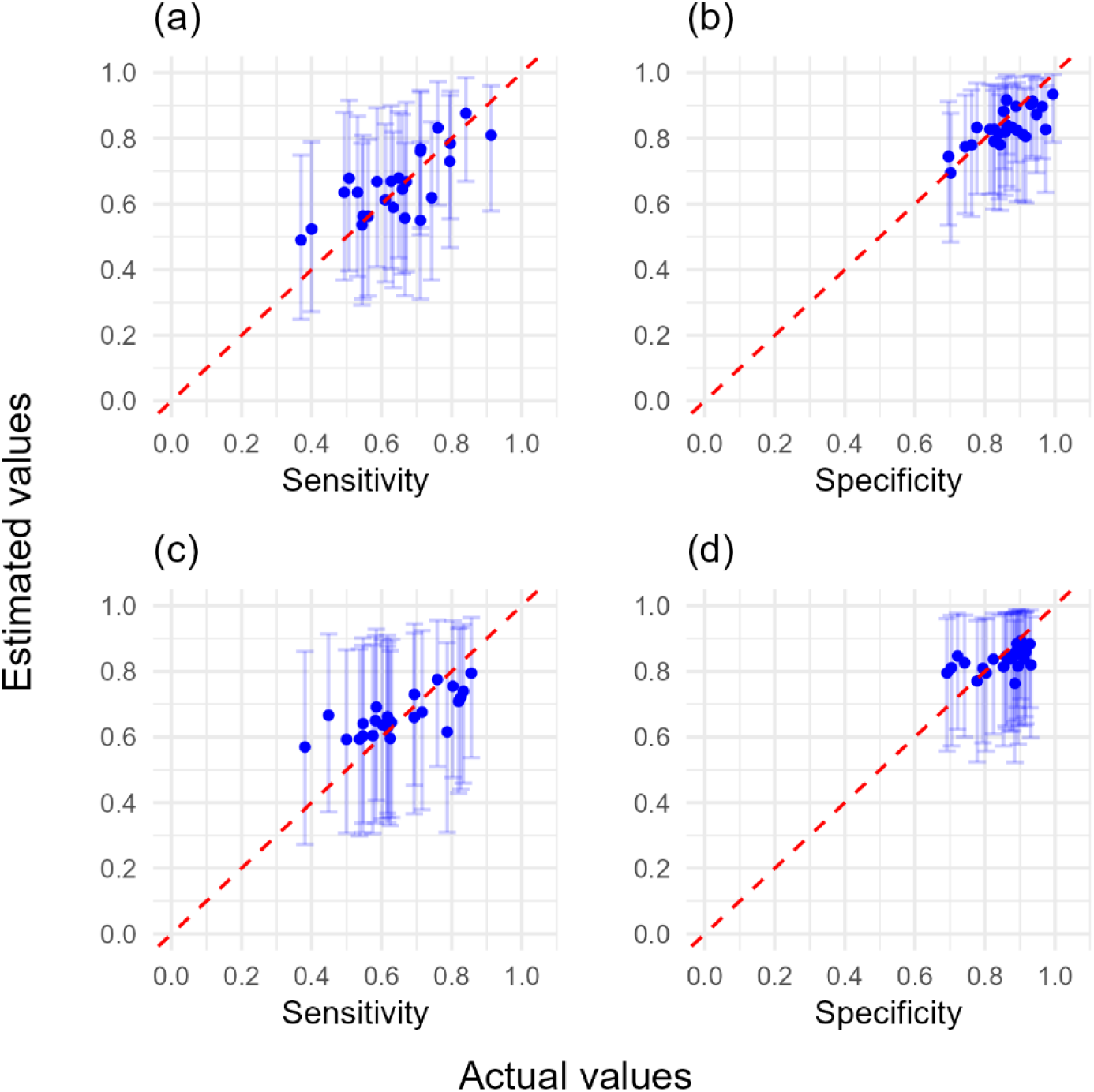
The workflow estimated sensitivity and specificity for simulated surveyors against actual sensitivity and specificity values (120 trees, good knowledge both symptoms). The workflow used good sensitivity and specificity prior distributions with (a, b) no covariance model with no simulated covariance, (c, d) covariance model with simulated covariance. In both instances 120 trees each were assessed in higher and lower true disease prevalence locations. Error bars represent 95% confidence intervals, and dashed line represents perfect agreement between workflow estimated and actual values.

**Fig S18.**
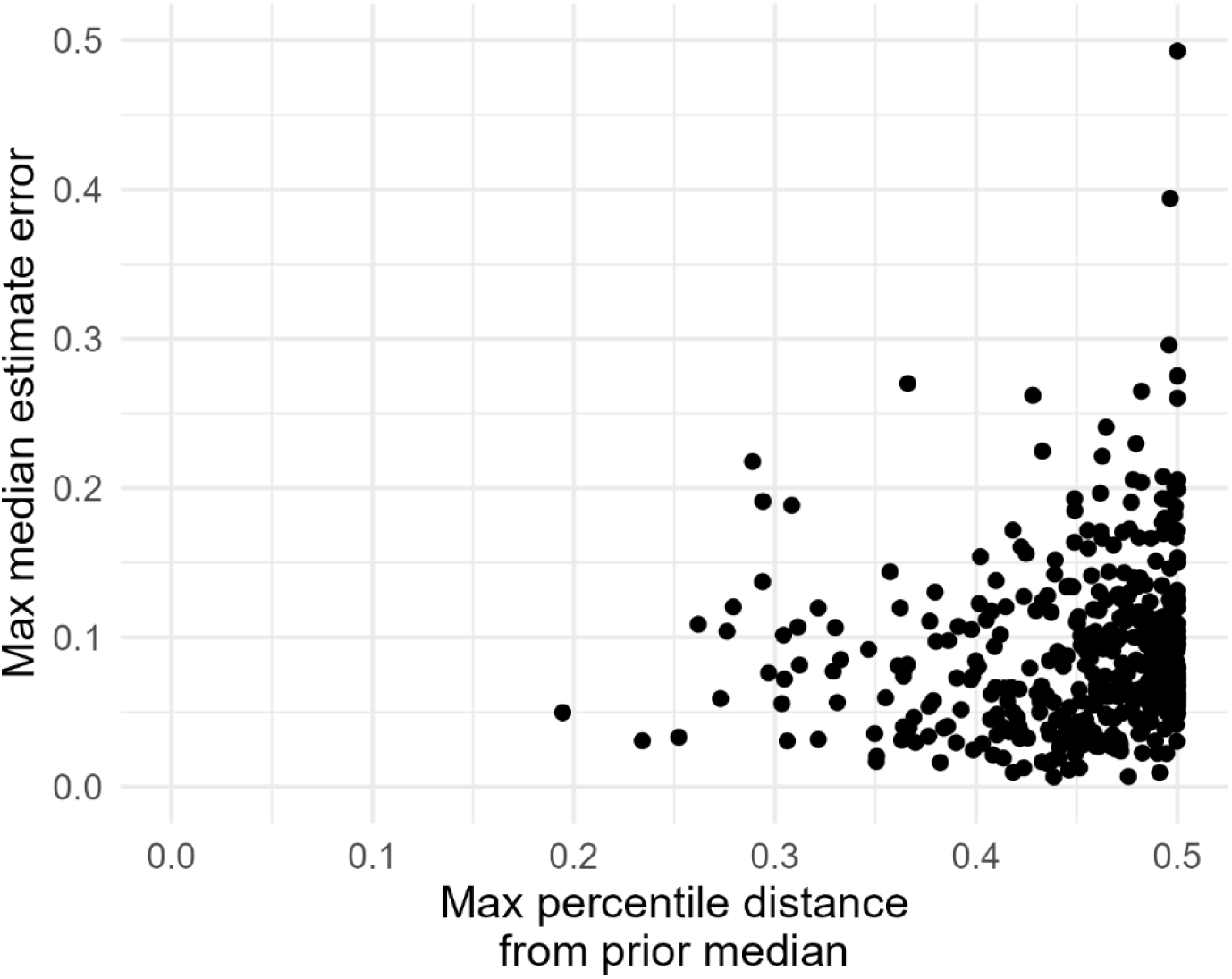
Workflow output errors in relation to the model prior distributions. The maximum error (i.e. difference from actual simulated value) of a median workflow estimate value for either the sensitivity or specificity against the maximum percentile distance of an actual simulated value from the median of a prior distribution (sensitivity, specificity, site true disease prevalences, or covariance parameters).

